# Asymmetric inheritance of actin regulatory machinery shapes cytoskeletal heterogeneity during early *Caenorhabditis elegans* embryogenesis

**DOI:** 10.64898/2026.03.22.713200

**Authors:** Grégoire Mathonnet, Roxane Benoit, Delphine Sunher, Nadine Arbogast, Elvire Guiot, Erwan Grandgirard, Anne-Cécile Reymann

## Abstract

Embryos experience challenges when transforming a passive oocyte into a polarized embryo. In *Caenorhabditis elegans,* polarisation is initiated at the one-cell stage and followed by a series of asymmetric or symmetric cell divisions. Patterns emerge via the regulated segregation of molecules at each cell division, including polarity proteins, cell fate determinants, transcription factors, mRNAs, and actin. The actomyosin cytoskeleton plays a crucial role in these early differentiation events, but how actin itself is segregated asymmetrically during the first divisions of the zygote remains poorly understood.

This study presents a thorough quantification of the spatiotemporal distribution of actin, the actin nucleators CYK-1 and Arp2/3 Complex, the capping protein CAP-1, and the E-cadherin HMR-1, from the zygote up to the 4-cell stage. To determine the potential for actin assembly of each early embryonic cell, we developed a novel assay combining *in vivo* microscopy-based quantitative analysis of the cytoskeleton, with *in vitro* actin polymerisation assays from single cell extracts named SCOPE-Single Cell cytOPlasm Extraction. SCOPE relies on UV laser ablation to empty blastomere cells into a polymerisation competent environment while following subsequent actin polymerisation. We find asymmetric segregations of most cytoskeleton proteins in favour of larger cells in a manner that is dependent on PAR polarity cues. AB and EMS cells being notably actively enriched compared to their sister cells, P cells being maintained in a state of low actin filaments. Interestingly, we discover an enrichment in ABp compared to ABa as division occurs, demonstrating that symmetric cell divisions are not exempt of actin enrichment. Additionally, this result illustrates that asymmetric distribution of actin-related proteins can precede known cell differentiation events. Taken together, these cytoskeleton heterogeneities are adding a layer to the complexity of cell fate acquisition mechanisms in the early embryo.

**Significance Statement:**

- The embryonic cytoskeleton can be actively inherited rather than passively distributed during early development.
- Actin assembly capacity can differ between blastomeres before fate diversification.
- Differential inheritance of actin regulatory machinery represents a potential mechanism linking cell polarity to developmental cell fate decisions.

## Introduction

*C. elegans* embryogenesis combines both an invariant lineage and early cell specification from the zygotic stage onwards. A combination of symmetry-breaking events of different nature enables the “tour de force” of assigning each cell its identity, thus determining the fate of the respective lineages in the first few divisions (Gönczy and Rose, 2005; Rose and Gönczy, 2014; Samandar Eweis and Plastino, 2020). Cellular polarisation of membrane-associated proteins, segregation of cell fate determinants, mechanical and chemical processes, non-uniform diffusion gradients, as well as phase separated granules or asymmetries in volume during division are a few examples of the required symmetry breaking events driving these cell differentiation events. The multiplicity of the factors involved is underlining the complexity of the scenario leading to the robust embryonic choreography at work throughout the *C. elegans* lineage.

The rapid segregation of some cellular compounds has been intensively studied during the zygotic anterior-posterior polarisation stage established rapidly after fertilization and leading to the first asymmetric division. During this phase, two simultaneous and coupled mechanisms are at work: on one side the distribution of partitioning defective (PAR) proteins into two membrane domains defining the Antero-Posterior (AP) axis, and on the other side the establishment of a cortical acto-myosin contractility gradient, inducing flows along the same axis (Rose and Gönczy, 2014; Samandar Eweis and Plastino, 2020). Cortical flow is dragging all actin-associated cortical proteins as well as membrane-associated proteins to the anterior side of the zygote, while flushing cytoplasm in the opposite direction. Advection due to frictional forces between the different layers (membrane, cortex and cytoplasm) but also chemical interactions between PAR proteins and some actin-binding proteins, notably modulating protein turnover in the cortex, are coupling flows to anterior-posterior polarity establishment (Delattre and Goehring, 2021; Goehring and Grill, 2013). Additionally PAR polarity at the membrane, via PAR-1 gradient kinase activity, defines cytoplasmic domains that are established most probably via differential molecular mobilities in the cytoplasm. Consequently, cytoplasmic concentration gradients are established in the zygote for different molecules including MEX-5, MEX-6, POS-1, PIE-1 or MEX-1 (Daniels et al., 2010; Griffin et al., 2011; Hoege and Hyman, 2013; Tenlen et al., 2008; Wu et al., 2015). The first asymmetric distribution of these cell fate determinants, between the larger AB daughter and smaller P1 daughter, is setting the stage for the following deterministic cell division pattern and invariant lineage of *C. elegans*.

In addition to PAR polarity proteins, studies have already revealed the anterior enrichment of some proteins present in the cell cortex. First of all, the actin filaments show asymmetries either using the MOE-1 labelling system (Shivas and Skop, 2012; Velarde et al., 2007) or using the Lifeact::mKate2 probe (Caroti et al., 2021; Reymann et al., 2016), even though these asymmetries are less prominent using the Lifeact::GFP or GFP::MOE probes (Hirani et al., 2019). The molecular motor, NMY-2, responsible for the contractility of the actin cortex (Pacquelet, 2017; Reymann et al., 2016; Scholze et al., 2018; Schonegg and Hyman, 2006) is highly asymmetrically in the cortex but is also showing gradients within the cytoplasm (Najafabadi et al., 2022), with locally restricted mobility in the anterior half (Tenlen et al., 2008). The scaffolding proteins, septins (Gilden and Krummel, 2010; Jordan et al., 2016; Nguyen et al., 2000), the E-cadherin HMR-1 implicated in adherens junctions (Padmanabhan et al., 2017), DYN-1, a protein that regulates endocytosis and actin comet formation (Ai and Skop, 2009; Shivas and Skop, 2012), as well as the Rho GTPase RHO-1, CDC-42 and their Guanine nucleotide Exchange Factor (GEF) ECT-2 (Scholze et al., 2018) do all also show some levels of asymmetries in the zygote. Regarding the essential formin in the early embryo, namely CYK-1, it has been reported that it is enriched in the extreme anterior cortex and then progressively switches to an increase in the extreme posterior cortex just before the anaphase cortical rotation (Zaatri et al., 2021). These results already point towards the presence of asymmetric distributions of actin-binding proteins as well.

In *C. elegans*, although the second division of AB, is symmetric and giving rise to ABa and ABp, these two blastomeres subsequently generate different lineages, the pharynx for ABa and the hypodermis for ABp, following a later induction by the P2 blastomere (Tax and Thomas, 1994). The third division is however asymmetric, in P1 cell a second PAR polarity gradient is established, leading to its asymmetric division into the larger ventral EMS cell and the smaller posterior P2 cell, from which will derive the germ cells lineage (Delattre and Goehring, 2021; Gan and Motegi, 2021; Gönczy and Rose, 2005; Horvitz and Herskowitz, 1992; Rose and Gönczy, 2014; Sulston et al., 1983). Due to egg shell confinement and imposed cell-cell contacts of the diamond-shaped 4-cell stage, a signal from P2 leads to the induction of EMS and ABp but not of its sister cell ABa, which has no direct contact with P2 (Evans et al., 1994; Rocheleau et al., 1997). Thus, *C. elegans* provides a unique model to investigate both asymmetric and symmetric cell divisions key for cell identity determination, which follow robust and highly stereotyped patterns within an exceptionally rapid developmental timeline—in less than an hour.

Taken together, different studies have made the link between asymmetries of mechanical properties of the acto-myosin cortex such as contractility or chirality and early embryonic *C. elegans* lineage (Naganathan et al., 2018; Pimpale et al., 2020; Pohl and Bao, 2010), but it has never been explored if some single cell differences exist in terms of actin-nucleating capacities after the first division which may impact cell identity acquisition. Here, we explore to which extent proteins responsible for actin nucleation show cytosolic asymmetries in the early embryo and follow their recruitment to the cortex. A key question is whether this machinery imposes limited availability in some cells, thus restricting locally actin polymerization capacities (Carlier, 1998; Suarez and Kovar, 2016). To address this question, we characterised the global dynamics of four main actin-binding proteins known to impact actin polymerisation and cortical mechanics from the 1- to the 4-cell stage (Naganathan et al., 2018). We chose two actin nucleators, the formin CYK-1 and the Arp2/3 complex via its subunit ARX-2, as well as a capping protein, CAP-1 and the *C. elegans* E-cadherin, HMR-1. CYK-1 belongs to the formin family, its human orthologue DIAPH1 is essential for the first cell cytokinesis completion (Jordan et al., 2016; Severson et al., 2002; Swan et al., 1998). The Arp2/3 complex is required for cortical filament nucleation and stability in early embryogenesis (Severson et al., 2002; Yan et al., 2022) and is mandatory for cell migration during gastrulation (Knight and Wood, 1998; Roh-Johnson et al., 2012; Roh-Johnson and Goldstein, 2009; Yan et al., 2022). Interestingly, balanced nucleation via Arp2/3 and CYK-1 set filaments homeostasis and are required notably for timely assembly and constriction of the contractile ring during cytokinesis (Chan et al., 2018) or filopodia nucleation (Hirani et al., 2019). The capping protein CAP-1 is also essential for proper actin organisation during the zygotic stage or in the adult germline by oppositely restricting polymerisation (Ray and Zaidel bar, 2021). Finally, HMR-1 is an important player to control cortical mechanics and as such is essential during diverse morphogenesis events, including blastomere polarity establishment or gastrulation (Caroti et al., 2021; Padmanabhan et al., 2017; Yamamoto et al., 2026). It is mainly involved in tethering the actin cortex to the cell membrane both at cell-cell contacts but also in non-junctional region where it additionally acts as a spatial regulator of Rho signalling (Padmanabhan et al., 2017).

To assess if actin polymerization capacity is equivalent in all cells of the early embryo and how actin cytoskeletal networks are inherited over the first few divisions, we performed a quantitative assessment of the availability of actin regulation machinery. To do so, we have set up a pipeline of live imaging spinning-disk microscopy and semi-automated quantitative image analysis, giving a reference map in space and time of endogenously expressed proteins densities in each cell as an output. This method allowed us to follow protein spatio-temporal dynamics at both cortical and cytoplasmic planes and compare the differential segregation between cells during the early dynamic steps of *C. elegans* embryogenesis. To expand these *in vivo* results and probe more directly actin content itself, we performed *in vitro* actin polymerization assays on single cell extracts using SCOPE-Single Cell cytOPlasm Extraction. This experiment allowed us to confirm differential actin content in sister cells. Therefore, our data provide a unique map of actin-related proteins in early embryos, with asymmetries correlated to differences in cell identity.

## Results

### Actin binding protein distribution over the course of the few first divisions

In order to follow the spatio-temporal distribution of primary regulators of the actin polymerisation in the early *C. elegans* embryo, we took advantage of CRISPR/Cas9 GFP knock-in strains (Table 1) for two actin nucleators, the CYK-1 formin and ARX-2, a subunit of the Arp2/3 complex, as well as the capping protein CAP-1 and the cadherin HMR-1. We focused our analyses from the zygote to the 4-cell stage (Figure 1A). Endogenous CRISPR labelling ensures that the fluorescence intensity acquired via high-resolution, high-sensitivity microscopy accurately reflects the level of endogenous protein: the intensity of an equatorial section of the embryo being proportional to the cytosolic concentration. At this localisation, cytoskeletal proteins mostly diffuse freely in the cell volume but can also interact with the filaments populating the cytoplasm (Raja Venkatesh et al., 2024; Velarde et al., 2007) (Figure 1B). The cytosolic concentration can thus be considered as a pool of proteins that undergo transition between activated and inactivated states. Quantification of overall cytoplasmic fluorescence intensity revealed differences in the mean intensity between labelled proteins (Figure S1A). For instance, endogenously expressed CYK-1::GFP and HMR-1::GFP show the lowest fluorescence level at this stages, ARX-2::GFP and CAP-1::GFP being clearly higher. This result revealed differences in the expression levels of these cytoskeletal proteins in the early embryo.

**Figure 1:**
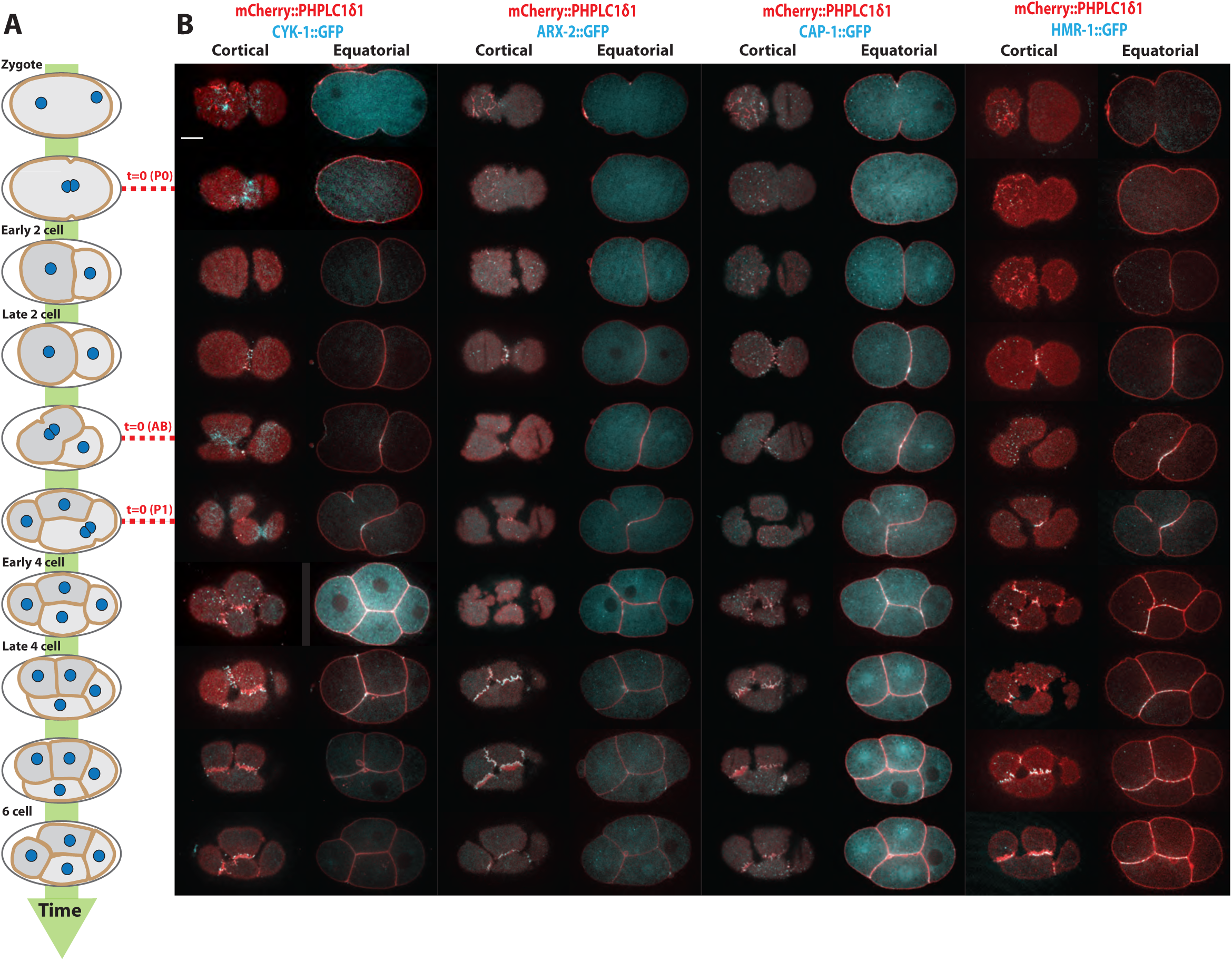
Distribution of some actin binding proteins over the first few divisions. (A) Schematic representation of the early steps of embryogenesis. The red dashed lines correspond to the cytokinesis onset of each cell stage used as time reference. (B) Spatio-temporal distribution of proteins of interest (cyan) together with the membrane marker (PHPLC1δ1, red), at the cortex (left column) and equatorial plane of the embryo (right column). Scale bar is 10 μm.

**Table 1:**
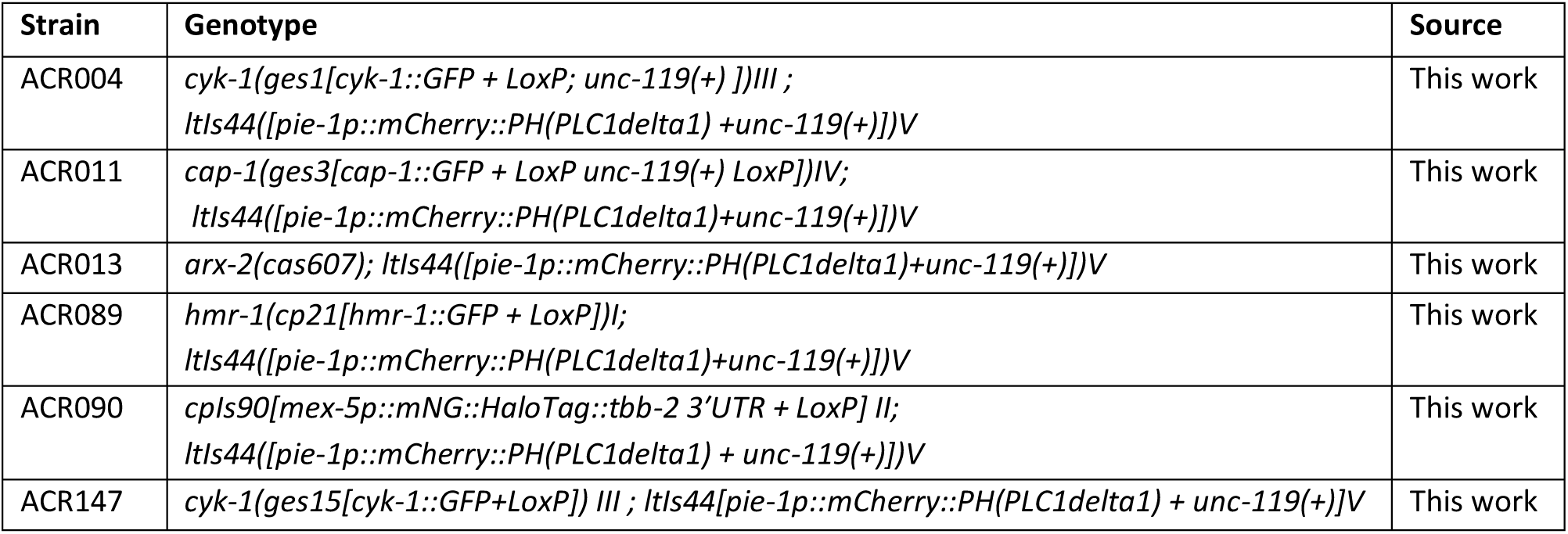
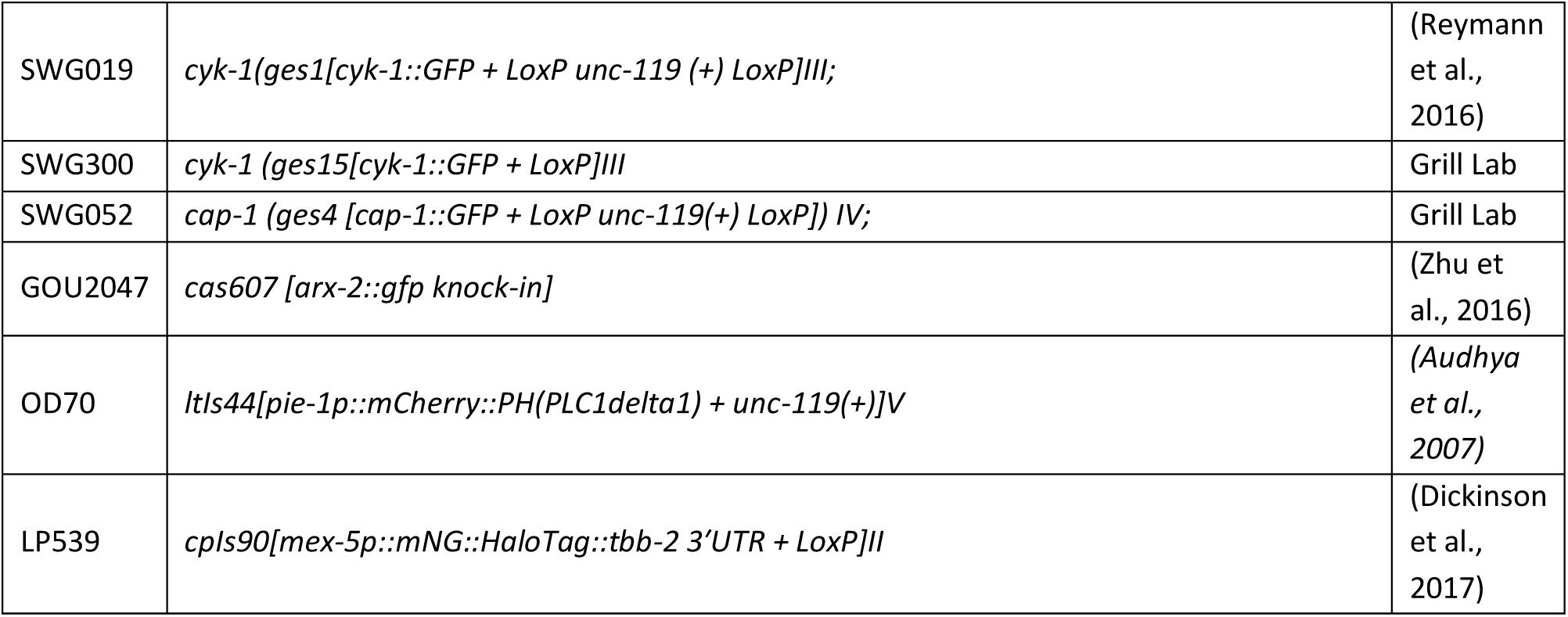
Newly generated strains for this study.

Acquisition of the fluorescent intensity at the cortical plane allowed to access protein density at the cell cortex, thus reflects the actively engaged proteins. As expected, additional shorter lived actin-rich architectures were abundantly observed in the cortical plane: CYK-1 rich cytokinesis rings appearing at each cell cycle, Arp2/3 rich endocytosis patches (Nakayama et al., 2009; Shivas and Skop, 2012), lamellipodial extensions at cell-cell contacts and filopodia that are clearly distinguished by their characteristic tips labelled with CYK-1 (Hirani et al., 2019)(Figure 1B).

### Cytosolic actin related proteins are differentially segregated between sister cells

To assess how actin related cytoskeletal proteins are inherited over the first few divisions in the early embryo, we followed the partitioning of the actin-binding proteins before and after each division step. We focused first on quantification of intensities in the cytoplasmic cross section of the embryo, thus on the global pool of proteins available. Single embryo quantifications of the time evolution of the mean intensity in each cell clearly showed some inequality of cytoplasmic mean intensities between sister cells (Figure 2A). For example, we observed that the mean cytoplasmic intensity in the larger AB cell is higher than in its smaller P1 sister cell for ARX-2::GFP and CAP-1::GFP but not very different for CYK-1::GFP. The ABp cytosolic fraction is enriched at the four-cell stage compared to ABa and EMS, and the smaller P2 shows significantly lower levels for all three proteins.

**Figure 2:**
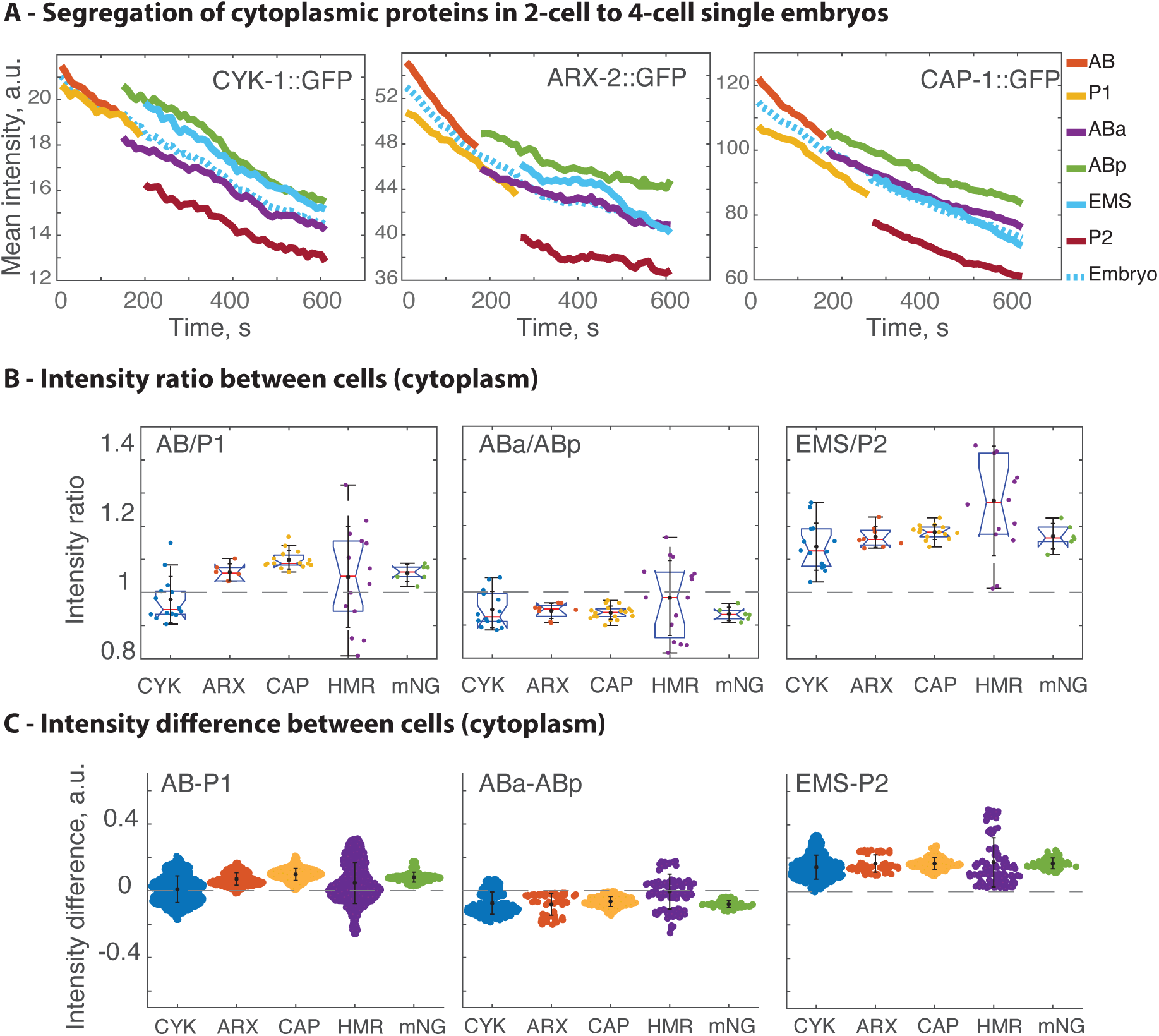
Cytosolic proteins are often asymmetrically partitioned between sister cells. (A) Time evolution of the mean intensity at the equatorial plane in each cell for a representative single embryo expressing fluorescently labelled ABP. Dashed lines represent the evolution of the mean intensity of the embryo (all cells). (B) Ratio of intensities for each cell pair measured in the cytoplasm and averaged during the whole cell cycle. (C) Differences of cytoplasmic intensities for each cell pair, calculated via the convolution of normalised density probability distributions taken at each time point and normalised by the mean embryo intensity for each strain. Individual data points, mean values and standard deviations are shown for B and C. The number of cells analysed for each category is given in the table Figure S1D.

To better assess cell-to-cell variability and thoroughly compare embryos, we used two different quantitative approaches. First, we calculated the ratio of the mean intensity between each cell pair taken at each time point, i.e. a proxy of fold change of concentrations between cells (Figure S1B). The time evolution in the cytosol showed a progressive increase of the ratio at the 2-cell stage but otherwise little variation (Figure S2B), we thus compared the averaged of these ratios over the duration of the whole cell cycle (Figure 2B, C, S3). The use of intensity ratio of two cells from the same embryo acquired at the same time point allowed to avoid the need of additional normalization between samples or strains and to correct for photobleaching. Second, we used the probability distribution function of the intensities found in each cell type to compile at each single time point, the absolute difference between each cell pair via the convolution of their probability distribution and normalised these differences by the overall levels of fluorescence intensity for each strain (Figure S1C). In both approaches, we added a non actin binding protein, the highly expressed cytosolic mNeonGreen (*mex-5*p::mNG::Halo transgene), for assessing passive cytosolic distribution.

Our two methods of quantification of fluorescent intensities, used as a proxy of protein differential concentration between embryonic cells, lead to similar results (Figure 2B, C, S3A). They showed an imbalance in favour of AB after the first division notably for CAP-1::GFP, ARX-2::GFP and the cytoplasmic mNG, whereas CYK-1::GFP and HMR-1::GFP distributions were more balanced between AB and P1. At the 4-cell stage, we observed a slight but constant bias in favour of ABp compared to ABa for all three actin binding proteins as well as for mNG, whereas HMR-1 remained more equilibrated. Stronger enrichments were observed for EMS relative to P2 for all five proteins. In conclusion, cytoplasmic differences visibly emerged from these first cell division patterns. Our results revealed more pronounced differences in cases of asymmetric divisions such as AB/P1 or EMS/P2, although some differences were also observed in the AB daughter cells. Thus, these observations challenge the view that AB division is perfectly symmetric before the induction due to contact with P2 (Gan and Motegi, 2021; Priess and Thomson, 1987). Indeed, we observed that differences between ABa and ABp were observed as soon as division occurred (Figure 2A). In conclusion, each blastomere between the 1- to 4-cell stages contains a different set of cytosolic concentration of cytoskeletal proteins, thereby contributing to acquire a different cell identity. Notably, the P1 and P2 cells were depleted of all protein content, presumably a hallmark of germ cells progenitors. Importantly, the presence of asymmetries in the segregation of cytosolic mNG proteins opens the possibility that asymmetries exist in terms of diffusion rates within the cytoplasm, irrespective of the nature of the protein (Hoege and Hyman, 2013). However, if some cytoskeletal proteins were segregated through passive diffusion, others such as CYK-1 and HMR-1, segregated more symmetrically, likely following a different segregation rule. To summarize, we conclude that specific patterns of partitioning of actin-related cytoplasmic components arise at these early stages. Notably, these patterns cannot be explained solely by a homogenous diffusion, thus distinct segregation mechanisms are possibly at work.

### Protein differential segregation is dependent on embryonic polarity

To further investigate the process by which segregation is controlled, we next performed a correlation analysis to assess in which conditions protein segregation deviates from a simple volumetric partitioning. The embryo initially divides while maintaining a constant total volume, successive division can be either asymmetric or symmetric, leading to major volumetric differences among cells (Cao and Wang, 1990). If segregation occurs passively: molecules partition proportionally to cell volume via diffusion within a homogeneous cytoplasm, both cells should inherit the same concentration, hence concentration ratio would be equal to 1 (Figure 3 A). Oppositely, in the case of a symmetric partitioning of molecules (quantities), concentrations vary according to the cell volume, thus concentration ratios vary as the inverse of the cell volume ratio (Figure 3A, blue dashed line). We used as proxy of cell-to-cell volume difference the ratio of the mid plane section area in our experiments. Our correlative plots of intensity over area ratios showed that CYK-1 and mNG intensity ratios spread horizontally, thus favouring the hypothesis of conservation of concentrations irrespective of cell volumes differences (Figure 3A). In contrast, ARX-2, CAP-1 and HMR-1 followed another trend, showing an increased cell-cell intensity ratio with increased volume difference. Such effect suggests an active amplification of asymmetries between cells, either via protein enrichment in larger cells or depletion in smaller cells.

**Figure 3:**
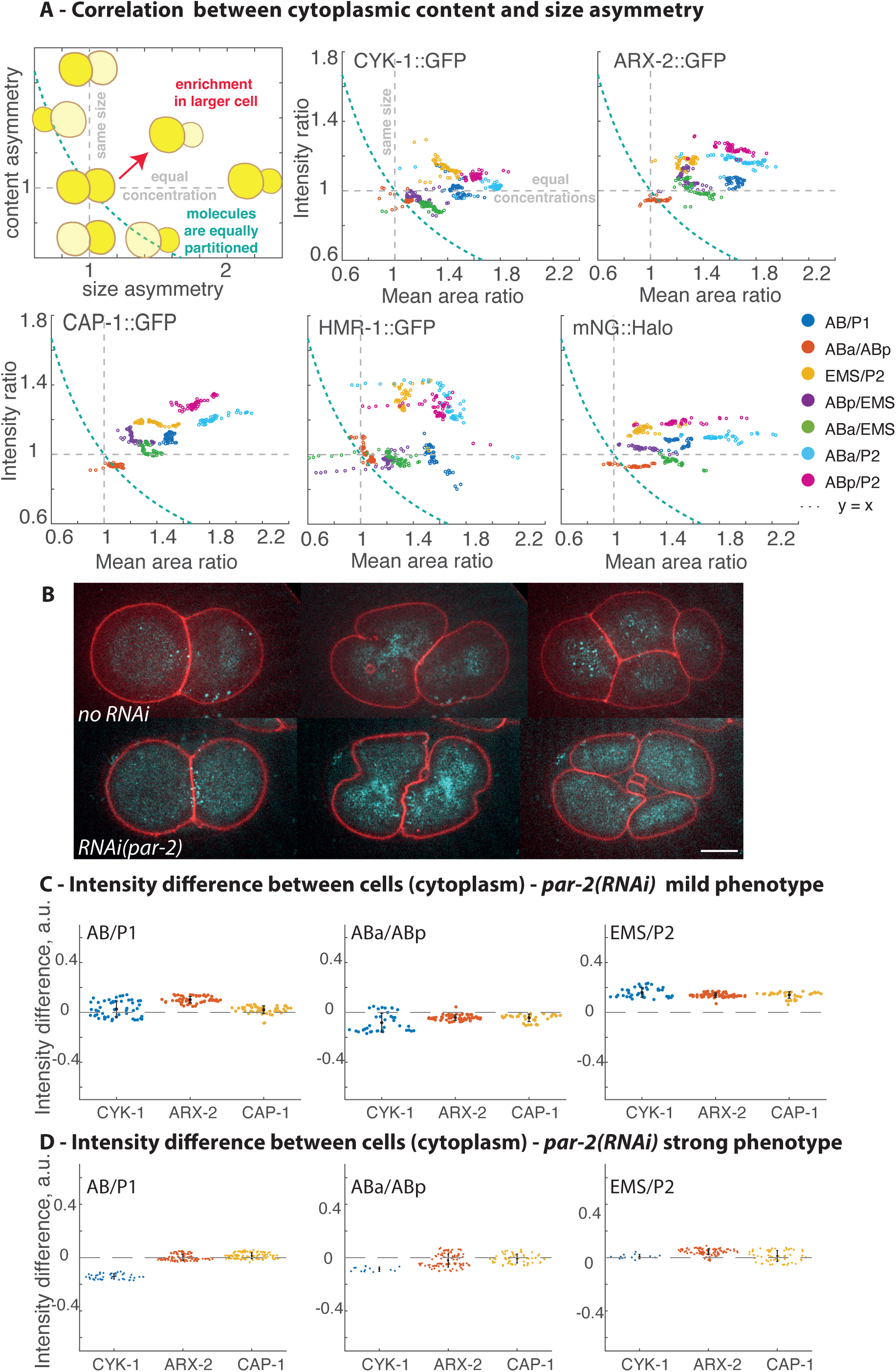
Perturbation of AP polarity impacts the distribution of actin binding proteins. (A) Illustrative phase diagram and correlation plots between mid-plane area ratio (used as a proxy of cell volume ratios) and cytoplasmic intensity ratio, shown for all cell pairs and proteins. (B) Representative images of embryos in absence and in presence of RNAi(*par-2*). Overlay of cortical CYK-1::GFP (cyan) together with the membrane marker in the mid plane section (PHPLC1δ1, red) are shown. Strong phenotypes are classified according to the abnormal 4-cell stage cell positioning (bottom right panel), while milder phenotypes maintain the normal cell-cell positioning as AB and P1 divided synchronously. In both RNAi conditions, the asymmetry in terms of volume is reduced for the cell pairs AB/P1 and EMS/P2. Scale bar is 10 μm. (C, D) Differences of intensities for each cell pair in the cytoplasm calculated via the convolution of normalised density probability distributions taken at each time point and normalised by the mean embryo intensity for each strain.

Note that these general trends of segregation are not applicable for all cell divisions or proteins. For instance, ARX-2 appeared in between the two general trends. Additionally, some cell pairs maintained a specific position irrespective of the protein assessed. Of note, ABa/ABp sister cells were systematically lying close to the stability point (1,1) with a slight shift towards the low right quadrant, thus very close to an equal partition of molecules but with some systematic enrichment in ABp. In contrast, the EMS/P2 asymmetric pair as well as all other cells compared to P2, lied systematically in the top right quadrant. Taken together, these analyses confirmed that differential mechanisms of protein segregation and homeostasis occur, depending both on the cell pair and on the protein considered. Several hypotheses could explain such difference. Protein synthesis is reduced in the early embryo but not fully shut down (Shukla et al., 2025), thus larger cells might increase the available protein concentrations via translation proportional to their size; whereas P cells already known to be transcriptionally repressed would follow the opposite tendency. Another possibility would be that differences in actin densities just prior division are associated to more binding sites in one half, thereby impacting the quantity of proteins inherited in each daughter cell as already suggested for myosin and cell fate determinants (Najafabadi et al., 2022; Tenlen et al., 2008).

We next assessed if the identified asymmetric distributions of cytoskeletal proteins were linked to anterior-posterior polarity establishment of the early embryo. We used RNAi of a polarity protein (*par-2)* to reduce the asymmetry of P0 and P1 divisions, thereby anteriorizing the embryo, synchronising the cell cycle at the 2-cell stage and leading to a disruption of cell positioning at the 4-cell stage (Beatty et al., 2013). Embryos subjected to *par-2(RNAi)* were classified in two categories according to their cell positions at the 4-cell stage: the loss of diamond shaped cell-cell contacts was classified as strong phenotype (Figure 3B), whereas normal cell-positioning was considered milder. In all cases, AB and P1 divisions happened synchronously, contrary to the WT condition. Quantifications were performed as previously described and revealed a clear reduction of the unequal partition of actin binding proteins, for all cell pairs (Figure 3C-D), except for CYK-1 that showed a peculiar loss of equal portioning between AB and P1. We conclude that actin related cytoskeletal protein segregation depends on embryonic polarity.

### Cortical abundance correlates with cytoplasmic abundance

Following these findings regarding cytoplasmic content segregation, we next investigated whether asymmetries in protein density also arose at the actin cortex where actin binding proteins are actively engaged. Note first that the cortical plane includes the actin cortex per se as well as many embedded actin architectures such as filopodia, endocytosis patches or lamellipodia. Second, we refer to the cortex as the cell surface area facing the eggshell i.e. the cell-free cortex. We excluded the cell-cell contact interfaces, notably due to the impossibility of attributing the proteins found at each of these interfaces to one or the other neighbouring cell. We thus compared the mean intensities at the cortical planes of each cell, following the same quantitative approach as for the cytoplasmic plane. Inequalities were observed, for instance the mean cortical intensity in the larger AB cell was higher than in its smaller P1 sister cell, for CAP-1 and ARX-2 (Figure 4A). The ABp, ABa and EMS cortical fraction were enriched at the four-cell stage compared to the total mean intensity of the embryo and to the P2 cell (Figure 4A, S3B). Note that time evolution of intensity ratios at the cortex can be more sensitive to cytoskeletal rearrangement, notably cytokinesis ring assembly, as observed for the 2-cell stage for CYK-1 (Figure S2B). Globally, we found similar results in the cortex to those quantified in the cytoplasm, both for intensity differences (Figure 4B, S3) and for the intensity ratio between each cell pair (Figure 4C, S3). AB was enriched in all proteins except CYK-1 compared to P1; EMS was enriched for all proteins relative to P2, with some more significatively than others; overall ABp was enriched for all proteins except for HMR-1. These results were confirmed by the correlation plots of intensities between the two different planes of observations (Figure 4D). In all cases, we observed a good linear correlation (Figure 4D). For CYK-1, the slope was close to 1 but higher for CAP-1 and ARX-2 (slightly below 1.3, Figure S3D). In the *par-2(RNAi)* condition, equalised cytoplasmic concentration correlated with an equalised cortical abundance (Figure 4E), except for CYK-1 again. Thus, cortical recruitment showed a strong correlation with the general availability of proteins in the cytoplasm at these stages. On one side, these results support the hypothesis that the cytoplasm constitutes a limited reservoir of available actin nucleators and capping proteins. This suggest that asymmetries in cytoplasmic concentration of proteins could affect the density of active proteins targeted to the actin cortical plane, thereby introducing cell specificity in the early embryo. On the other side, it is possible that asymmetrical recruitment at the cortex of actin binding proteins impact their cytoplasmic mobility, thereby affecting their segregation during division.

**Figure 4:**
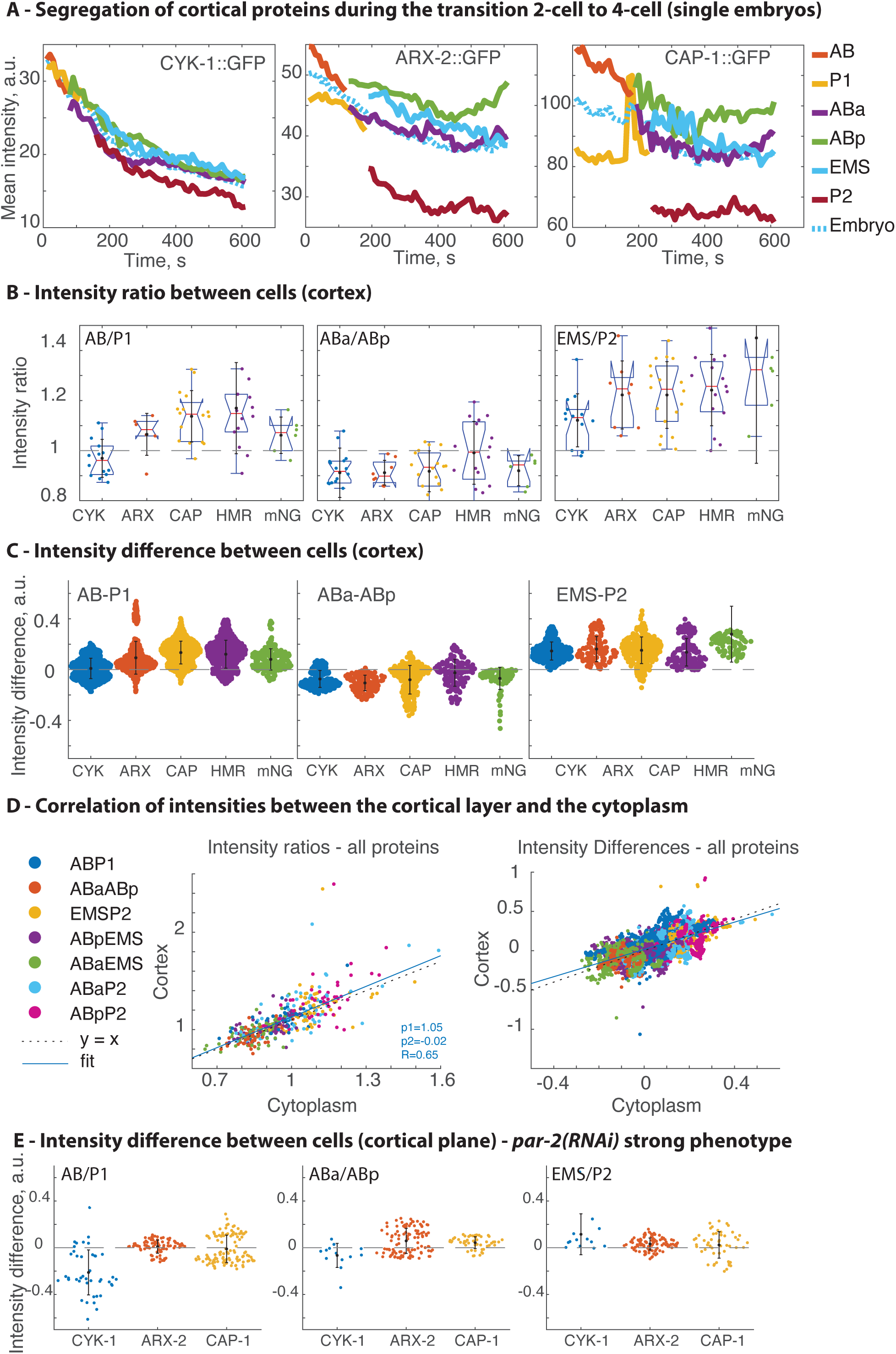
Protein densities show enhance differences at the cortex compared to the cytoplasm. (A) Time evolution of the mean intensity at the cortical plane in each cell for a representative single embryo expressing GFP-labelled ABP. Dashed lines represent the evolution of the mean intensity of the embryo (all cells). (B) Ratio of intensities for each cell pair measured in the cytoplasm and averaged during the whole cell cycle. (C) Differences of intensities for each cell pair at the cortex, calculated via the convolution of normalised density probability distributions taken at each time point and normalised by the mean embryo intensity for each strain. Individual data points, mean values and standard deviations are shown for B and C. (D) Correlation plot of mean intensities in the cortex and in the cytoplasm (all cells and all proteins). Linear model from the fit is shown as continuous line (f(x) = p1*x+p2) and f(x)=x is shown as dashed lines. See Figure S3 for individual plots. (E) Differences of intensities for each cell pair in the absence of presence of RNAi(*par-2)*.

### Novel method to probe single cell actin content

Beyond evaluating the differential abundance of actin-binding proteins, we further investigated whether embryonic cells might also exhibit variability in their actin content. Assessing precisely actin monomers or filamentous concentration within a particular cell is a complex task which relies on indirect measurements and technical tricks (Cramer et al., 2002; Gonzalez Rodriguez et al., 2023; Kiuchi et al., 2007; Koestler et al., 2009; Lee et al., 2013; Malla et al., 2022; Skruber et al., 2018; Vitriol et al., 2015). Indirect labelling via phalloidin or Lifeact probes is possible but can lead to different results depending on the probe used and normalisation process (Caroti et al., 2021; Courtemanche et al., 2016; Hirani et al., 2019; Motegi and Sugimoto, 2006; Reymann et al., 2016; Skruber et al., 2018; Yamamoto et al., 2026). We thus decided to develop an alternative method to assess actin polymerisation in a cell-specific manner that is suitable to compare the few individual cells of the early *C. elegans* embryo, that we named SCOPE, for Single Cell cytOPlasm Extraction. We used *in vitro* actin polymerization assays on single cell extracts to quantitatively evaluate both F-actin amount and the overall actin polymerization capacities, defined as the mass of polymerised actin that each cell is capable to form in a standard medium. Targeted UV laser ablation was used to perforate individual cells, leading to the extrusion of most of their cytoplasmic contents and the concomitant fragmentation of their actin networks (Figure 5A, S5). In the appropriate buffer condition containing Phalloidin-Alexa-488, we then followed the rapid actin polymerization outside of the cell ghost (Figure 5A, S5). Phalloidin, which binds to the side of filaments, labels both filaments assembled in the embryo prior ablation and those actively polymerizing in the cell extract. Oppositely, supplementation with low amount of purified monomeric actin coupled to Alexa-568, revealed only the filaments actively polymerizing after ablation- as the supplementation of G-actin is in the range of the critical concentration of actin bellow which spontaneous polymerization occurs (Figure 5A). Dual labelling confirmed that actin seeds are extracted from the lysed cell (green fragments) and rapidly elongate together with newly formed filaments, before reaching an equilibrium state when all G-actin is consumed (yellow fragments, Figure 5A, S5). Additionally, to verify that active polymerization is indeed at the origin of the filaments observed, a control condition was performed with Cytochalasin B, a drug inhibiting actin polymerization. This resulted only in short filaments observed in close proximity to the lysed cell (Figure 5A). To quantify which percentage of cellular actin is extruded and effectively released in the extract, experiments were conducted in presence or absence of detergent. As expected, addition of Triton Tx-100 to the medium lead to membrane disassembly, allowing the release of an increased amount of cellular content and increased the number of filaments observed (Figure 5A).

**Figure 5:**
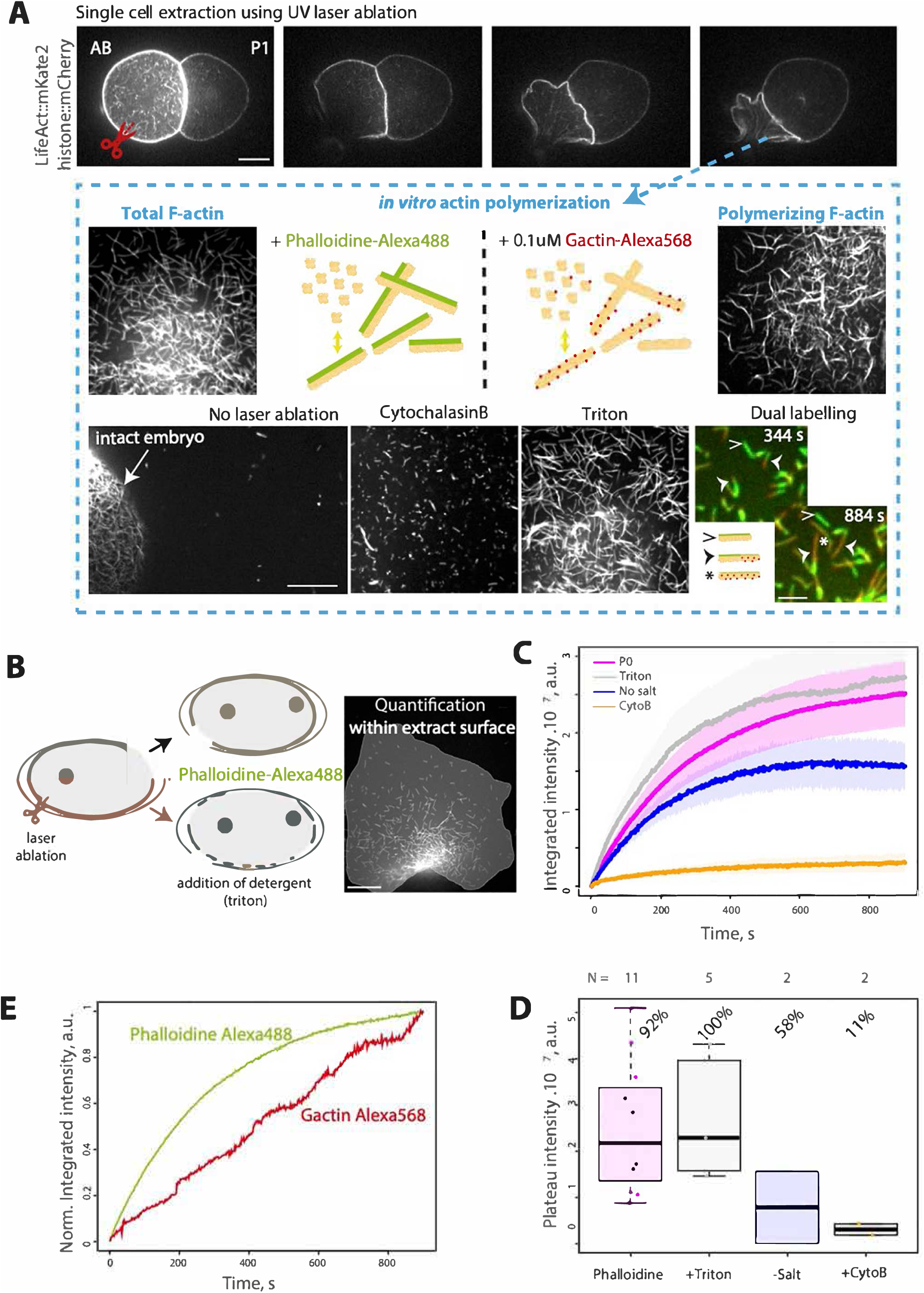
(A) Representative images of a single cell extraction using UV laser ablation, image of the cell prior ablation and just after the induced local cut (red scissors). Representative images of *in vitro* polymerisation from such single cell extracts after 15min in the different conditions. (B) Illustration of the two processes cell lysis via laser ablation with or without detergent. Image of the region of interest used for quantification purposes. (C) Time evolution of the integrated intensity profile of P0 cells in different experimental conditions and values obtained in the plateau region (D) together with the calculated percentage taken the +triton condition as reference. (E) Time evolution of the integrated intensity profile in the phalloidin and G-actin conditions.

During polymerization, we measured the fluorescence intensity in the area occupied by the extract. This integrated intensity rapidly reached a plateau value (Figure 5B, C). As in standard actin pyrene assays, the intensity at the plateau scales with the F-actin content at steady state. Using the mean intensity of this plateau we can estimate that experiments without triton represent 94% of the cellular content released in the presence of triton, thus supporting that a large amount of the actin cell content is released by UV ablation (Figure 5D). Applying the same quantification method, the presence of the Cytochalasin B led to a strong reduction of the intensity measured with Phalloidin labelling (14% F-actin remaining). Note that we observed a limited diffusion of filaments around the embryo debris (Figure S5A), and constraining the experiment in a smaller volume via PDMS microwells did not yield different results. Thus, to maintain reasonable experimental throughput, we decided to simply restrain the quantification to the visible extract area containing the majority of filaments (Figure 5B). To conclude, our experimental design allows to distinguish polymerized and actively polymerizing filaments from a single cell extract bellow the nL volume range, thus quantitatively assess single cell total actin content.

### Cell specificities are revealed in terms of actin polymerization capacity

This experimental design was used to compare single cell actin polymerization capacity from 1-cell to 4-cell stages (Figure 6A). We quantified the total amount of released actin using Phalloidin labelling from single cell extracts. Measurements of the integrated fluorescence intensity over time for the different extracts increased and reached rapidly a plateau (Figure 6B), allowing us to estimate the total actin content extruded out of these cells following laser ablation (Figure 6C). After 15 min, the F-actin networks obtained for the different cell extracts showed significant differences. P0 cell extracts exhibited the highest actin polymerisation capacity, consistent with the larger size of the zygote. Polymerisation capacities were similar between AB and P1 cell extracts. At the 4-cell stage, P2 cell extracts exhibited the lowest actin polymerisation capacity. This result is correlating with our *in vivo* observations that showed lower concentrations of actin nucleating proteins found in P2.

**Figure 6:**
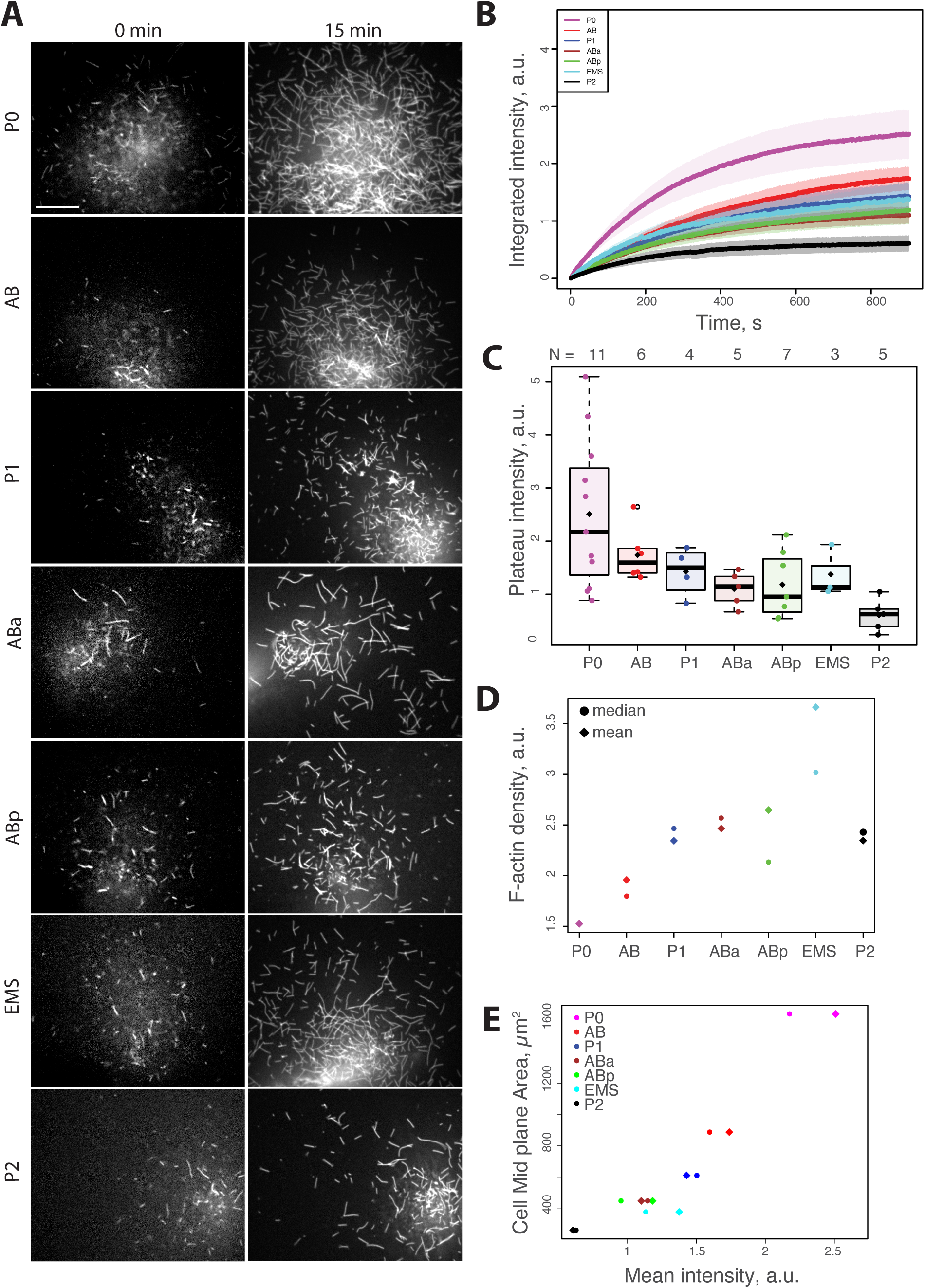
(A) Representative images of *in vitro* polymerisation from single cell extracts just after ablation and after 15min of polymerisation. Scale bar is 10 μm. (B) Time evolution of the integrated intensity profile of these cells. (C) Intensity values obtained in the plateau region. N is the number of cells analysed for each category. (D) F-actin density is calculated using the plateau intensity value normalised by the midplane area of the corresponding cells, mean (diamond) and median (circle) are shown. (E) Correlation between mean intensity values (plateau) and midplane area of the corresponding cells.

To estimate the differences in terms of actin concentration available for polymerisation within their cell of origin, we additionally normalised these values using either our measured midplane cell area prior ablation or published values of these cell volumes (Figure S5D, E). Surprisingly in terms of these normalised actin content (Figure 6D, S5E), P0 fell at the lowest position, P1 was over AB and EMS showed the highest values (Figure 6E). Thus, smaller cells such as P1 or P2 have lower total actin content but maintain surprisingly a high actin concentration. This observation may reflect the progressive increase in surface-to-volume ratio during embryonic cleavages, which imposes a greater demand for cortical actin assembly. Additionally, EMS larger estimates of actin concentration might be linked to its assembly of large lamellipodial protrusions extending over its neighbouring ABa and ABp cells. Altogether, first these results indicate that there is not a strict correlation between cell volume and the estimated actin content available for active polymerisation, notably for larger and smaller embryonic cells (Figure 6E). Second, it confirms that there is a fine tuning of the availability of polymerizable actin in each of these blastomeres in addition to differences in the availability of actin binding proteins.

## Discussion

The early embryo relies initially mainly on maternally provided proteins loaded in its large oocyte (Zacharias and Murray, 2016). In *C. elegans*, this initial pool of proteins is split through divisions into the different daughter cells, asymmetrically or symmetrically, along a fixed lineage and must be sufficient to ensure the success of these first initial divisions (Rose and Gönczy, 2014). Asymmetries have already been characterised for several proteins, including cell fate determinants and differences in cell contents were shown to be drivers in the process of early cell identity acquisition. Regarding cytoskeleton proteins, only few studies mentioned asymmetries between cells, as for actin (Reymann et al., 2016; Shivas and Skop, 2012) or myosin (Munro et al., 2004; Pacquelet, 2017; Reymann et al., 2016; Schonegg and Hyman, 2006), for which a gradient of free NMY-2 molecules in the cytoplasm was recently characterised (Najafabadi et al., 2022). Our findings demonstrate that the temporal and spatial unequal partitioning of key actin assembly regulators—including a formin, the Arp2/3 complex, capping proteins, and the cadherin HMR-1—occurs in both in the cytoplasm and at the cortical plane during cell division. Concentration gradients of freely diffusing actin-binding proteins can form within the cytoplasm, analogous to those previously described for cell fate determinants such as PAR proteins and MEX-5 (Daniels et al., 2010; Hoege and Hyman, 2013; Tenlen et al., 2008). For example, AB and EMS cells inheriting more actin related proteins than their sister cells P1 and P2. Importantly, we show that these asymmetries in actin-related content depend on polarity cues. We can thus conclude that polarity determines how many actin-related proteins segregate in each polarity domain. Together our work paints the picture whereby controlled adjustment of cortical and cytoplasmic actin amounts prior to cell division determine how many actin proteins is inherited.

A surprising result from our work is the precocious asymmetry observed in AB daughter cells, ABa and ABp. Concentrations of CYK-1, ARX-2 and CAP-1 are spatially enriched in the ABp cell from AB division onwards (Figure 2A, 3A), thus indicating that a concentration gradient is probably set ahead or concomitant to AB division. Interestingly this difference is not observed with HMR-1::GFP, suggesting that it is specific to the cortical actin network factors. This difference arises prior to the known Notch signalling event, in which direct contact between P2 and ABp cell induces ABp cell fate specification (Priess, 2005). In other words, asymmetric distribution of actin-related proteins can precede cell fate acquisition in this cellular context. Further perturbation experiments using partial RNAi depletion or mutants will have to be performed to decipher what is the mechanism at work accounting for these differences. It is also very probable that different mechanisms act together to ensure the robustness of development- as is already described for polarity establishment in the zygote.

Along the lineage the increase of surface-to-volume ratio is imposing a continuous increase of cortical surface coverage by actin filaments. Such necessity might explain the general increase in the content of actin binding proteins observed in our *in vivo* quantification between the 2-cell and 4-cell stage (Figure S3D). The P2 cell however stood out from the other cells, showing both a significative depletion of proteins as well as presented an important reduction in the total amount of extruded actin -even though when normalised to cell volume this difference seems reduced. This raises the possibility that the P2 cell, and by extension the whole P lineage, is maintained in a state of low actin filament state. Oppositely EMS cells, in addition of showing high levels of ABPs, showed a high actin content notably when normalised by cell volume. These observations do correlate well with the capacity of EMS to build large lamellipodia extensions migrating over it neighbouring cells.

Asymmetries in the inherited distributions of cytoskeletal proteins are likely to amplify cell to cell differences in terms of actin assembly dynamics along the lineage of the early *C. elegans* embryo. Previous measurements focusing on CYK-1 single particle tracking to assess actin filament elongation rate between the 1-cell to 4-cell stage showed no significant differences in terms of elongation rate, suggesting no difference in the availability of actin monomers within embryos (Costache et al., 2022). An alternative hypothesis would be that filament growth is limited by another factor, such as the dissociation of profilin from actin monomers during filament incorporation in presence of high actin concentrations (Funk et al., 2019), thereby limiting the consequence of changes in the availability of actin. Modulating the availability of actin nucleators could be an alternative mechanism to modify differentially actin polymerisation capacities between cells. Additionally, nucleators’ availability impacts the physical properties of their cortex. For example, the hydrodynamic length is reduced by CYK-1, ARX-2 or CAP-1 depletion, leading to a decrease of cortical chiral flows in the zygote (Naganathan et al., 2018). More generally, the balance between actin nucleation via formin or Arp2/3 is controlling the density and degree of entanglement of actin networks, and the rate of filament capping is key to set filaments length. Note that all such changes in actin binding protein activity also impact the amount of monomeric actin available and the ratio between monomers and filaments, thus providing feedback loops impacting actin homeostasis. To conclude, some tight regulation of concentration gradients of proteins such as CYK-1, ARX-2 and CAP-1 is important to define cell specificities both in terms of cytoskeletal architectures and dynamics scaling up to specificities in terms of mechanical properties. Going a step further, as actin cytoskeleton and cell mechanics are intrinsically linked to cell identity, we can hypothesise that these differences are possibly playing a role in the cell potency during these early embryonic stages. Thus, differential inheritance of actin regulatory machinery represents a potential mechanism linking cell polarity to developmental cell fate decisions.

We found that only CYK-1 and the diffusing mNG seem closer to a passive segregation - following a volumetric partitioning of equal concentrations. In most cases, actin binding proteins did follow a ratio strongly in favour of larger cells, indicating that there might be an active mechanism at work to regulate the extend of asymmetric segregation. Our findings do not allow us to distinguish the causal relationship between actin related cytoplasmic and cortical asymmetries. One hypothesis would be that cortical regions with higher actin densities, such as the anterior half of the zygote, by offering more binding sites for actin binding proteins could maintain local enrichments in the neighbouring cytoplasm. Feedback mechanism might amplify these gradients over time: increase of local concentration of nucleators leading to increase in filaments thus more binding sites for ABPs. The binding and unbinding rates to actin filaments as well as overall turnover of actin networks would thus be key parameters controlling such mechanism. Observation of the distribution of the two Lifeact probes in the early lineage could support this hypothesis. Lifeact is a peptide binding to actin filament with fast binding and unbinding kinetics, it was previously shown that when coupled to mKate2 fluorescent probe its turnover was one order magnitude slower than when bound to GFP (Hirani et al., 2019). As a result Lifeact:mKate2 turnover is comparable to the turnover of cortical F-actin in the *C. elegans* cortex (Robin et al., 2014). In the early embryo, we have observed an enhanced asymmetry in Lifeact::mKate2 compared to Lifeact::GFP. It is notably enriched in the anterior side of the zygote up to the 4-cell stage (higher in AB at the 2-cell stage and more intense in Aba/ABp compared to EMS and P2). Our present quantitative analysis of actin related content is pointing towards the presence of an asymmetrical distribution along the same direction than this actin binding probe. In addition, our *in vitro* polymerisation assay also clearly points towards large differences in the total actin content between cells, even though it is less clear if these translate to effective differences in terms of available actin concentrations. In the future, a direct actin labelling method would be required to quantify with higher precision changes in actin densities between cells. To conclude, a minor anterior enrichment of cortical density in the zygote can be sufficient to maintain cellular asymmetry of a binding partner, which would be released upon disassembly of actin only. For this reason, we predict that the turnover rate of actin binding proteins might be key in the process of asymmetric inheritance in the lineage. Note that in *C. elegans* early embryo, actin turnover was estimated in the same range as CYK-1 formin bulk cortical turnover rate (Costache et al., 2022), thus this prediction could be valid, as well as for the Arp2/3 or Capping Protein which are thought to be longer-lived on actin filaments.

Of importance as well, in the cell divisions studied, the transgenic highly expressed free fluorophore mNG was observed as not strictly partitioning equally at each cell division, showing an enrichment notably for AB and EMS. This has never been reported so far and might be a hint that gradients in overall protein mobility within the cytoplasm might exist and slightly influence protein concentrations in these large cells. However, some singularities to this general trend were also observed for some actin binding proteins, indicating that some specific mechanisms may also exist, either to maintain equality between the cells or oppositely to enhance a spatial distribution pattern. For instance, the CYK-1 formin is surprisingly equally distributed between AB and P1, and this equality is lost in the *par-2(RNAi)* condition. This is not the case for the ARX-2 or CAP-1 which maintain more asymmetries overall in the cortex and cytoplasm before and after division. An alternative explanation for mNG asymmetry could reside in asymmetric distribution of organelles within the cytoplasm modulating via exclusion the quantity of diffusive proteins. In addition, general cytoplasmic streaming in opposite direction to cortical flows could also play a role in the concentration gradient observed for these two proteins.

In conclusion, we have established a spatio-temporal map of the early *C. elegans* embryo with respect to actin and key actin binding proteins at a high resolution. Our work raises the question of a potential embryonic control on differential cellular actin homeostasis and actin cytoskeletal steady states. We favour the hypothesis that concentration gradients could be sufficient to drive and maintain cellular asymmetries both in the cytoplasm and in the cortex for some important actin binding proteins. Some of the acquired cell specific cytoskeleton properties might be instrumental in the subsequent steps of cell differentiation events in the early *C. elegans* embryo. This knowledge will be instrumental in future studies aiming at understanding how actin directly impacts cell differentiation processes.

### Limitation of the study

Although we performed rigorous quantitative assessments of single cell contents, our study is subject to certain limitations that warrant consideration. First, due to a low thickness of the cell cortex we cannot exclude that fluorescence from the cytosol is impacting the fluorescence in the cortex, notably in the case of low density of protein recruitment in the cortex. This is especially the case for control mNG::HALO that does not attach specifically to the actin cortex. Similarly, as noted in the results section, our quantification focused on the cortex facing the eggshell, rather than on cell–cell contact regions, which are highly enriched in actin filaments. Another limitation comes from the fact that we cannot distinguish between active versus inactive proteins, notably for nucleators. Quantification of the availability of some key proteins required upstream in the signalling cascade would be complementary to our study to decipher the limiting factors. Finally, translating fluorescence intensity into accurate estimates of cellular concentrations represents significant challenges (Aroush et al., 2017; Gonzalez Rodriguez et al., 2023; Kiuchi et al., 2007; Koestler et al., 2009; Malla et al., 2022; Vitriol et al., 2015) and many additional parameters need to be considered while doing so. Notably, the cytosolic volume—distinct from the total cell volume due to organelles—can influence concentration estimates (Rodriguez et al., 2023), especially in P-lineage cells enriched in P-granules. For this reason, the use of cell volume as a mean to normalise the released actin content to compare possible actin concentration between cells should be taken with precaution.

## Materials and Methods

### Strains maintenance

Animals were cultured at 20°C on Nematode Growth Medium plates seeded with OP50 (Brenner, 1974) at 20°C. The strains used in this study are presented in the following table.

### Microscopy

L4 adults were collected on separate plates 24 hours before embryo imaging. Hermaphrodites were dissected on coverslip in M9 buffer and mounted under 2% agarose pad. Coverslip was sealed to slide with melted VALAP (1:1:1, Vaseline, lanolin and paraffin wax).

All microscopy samples were collected using an Inverted Nikon Eclipse Ti configured by Gataca Systems (Massy, France) and equipped with a Yokogawa CSU-X1 scan head, a simultaneous dual camera with two Prime 95B cameras (Photometrics) and a 100×1.4 NA objective lens. Simultaneous acquisition in dual colours was performed using 491nm and 561nm laser lines with a combination of emission filters for each camera (dual band 506-548/620.5-669.5 and 609/54) after splitting with a dichroic at 560 nm. Image acquisition was controlled by MetaMorph software (Molecular Devices, Sunnyvale, CA), in a temperature-controlled room set to 20°C. Imaging conditions were set to assure minimal photobleaching and no photo-toxicity in order to allow for live observation in 3 optical sections over 10 minutes of embryonic development but with sufficient sensitivity enabling the detection of low expressed levels of proteins (200ms exposure time with 20% power of 491nm laser and 30% for 561nm laser). Each Z-stack is timely separated by 10s for a total duration of 10min and maintained in focus using the Perfect Focus controller. Thus, each timepoint is composed of 3 different confocal planes spaced by 3µm, the cortical plane (Z=1), the intermediate plane (Z=2), the equatorial plane (Z=3).

### *In vitro* cell extracts

L4 adults were collected 24 hours before embryo imaging. Hermaphrodites were dissected on a cleaned coverslip coated with BSA 5% in 5µL of actin polymerization buffer (15 mM imidazole, pH 7.0, 74 mM KCl, 1.5 mM MgCl2, 165 mM DTT, 2 mM ATP, 50 mM CaCl2, 5 mM glucose, 30 mg/mL catalase, 155 mg/mL glucose oxidase, 0.75% methylcellulose) either supplemented with Phalloidin-488nm (1:40 000 diluted in G-buffer; A12379) and/or 0,1µM of purified G-actin-Alexa568 (from rabbit, gift from Christophe Guerin, Laurent Blanchoin). For Cytochalasin B experiments, 1µM (diluted in DMSO; C6762) has been added to the actin polymerization buffer. For triton experiments, 0,1% of Triton TX100 has been added to the actin polymerization buffer. Note that is not used to compare single cell contents to avoid contamination from neighbouring cells. Coverslip was sealed to slide with VALAP (1:1:1, Vaseline, lanolin and paraffin wax). Imaging was performed on the same microscope as described above; in single camera mode (filters: 525/50 (GFP) and 605/64 (RFP)) and using a different objective (Leica HC Plan Apo 100x/1.4-0.7oil WD 0.1mm). Cells of interest were lysed using UV laser ablation (iLas2 FRAP module from Gataca system equipped with a 355 nm, 0.6 ns pulsed, 5W laser (Teem Photonics)). Images were acquired prior ablation (notably a mid-plane section in phase contrast) and for 15 minutes after ablation in the plane adjacent to the coverslip with a framerate of 2s (30% laser power and 200ms exposure time). Focus was maintained using the Perfect Focus controller.

### Image analysis (*in vivo*)

Cell outline was segmented using an interactive segmentation tool based on machine learning algorithms, ILASTIK (Berg et al., 2019). The training was performed on the equatorial (Z3) and the intermediate plane (Z2). The segmentation of the complete timelapse was then obtained automatically. As the Z1 plane was more prone to segmentation errors, we preferred to use the clean Z2 segmented regions adequately cropped to fit the cortical area in the Z1 plane.

In order to realign each embryo over its developmental time, we used a reference time point for each cell cycle - chosen as the onset of membrane ingression observed in the equatorial section (Z3) during cytokinesis onset (Chan et al., 2018) (Figure 1A).

We quantified the average fluorescence intensity for each cell at each time point and for each optical section using Matlab. Prior analysis, we corrected for inhomogeneity of the field of view. We used two approaches to compare intensities between cells. First, we performed cell to cell ratio for each time point either for cytosolic or cortical mean fluorescence intensities. This mean intensity ratio allowed us to estimate the relative proportion of fluorescently labelled proteins between cells. Second, we used the probability distribution function of the intensities found in each cell. We compiled the convolution of the normalized distributions for each cell pairs giving in a statistically significant manner the cell pair content differences for all time points (Figure S1). In order to normalize intensity differences between strains showing significative differences in the level of expression of the protein of interest, we normalized these differences using the embryo mean intensity taken for all samples and all frames. Correlation plots were performed, using the mean value of each cell to cell mean intensities ratio at the cortex and at the cytoplasm, fitting a linear model (f(x) = p1*x+p2) while excluding the outlier values with cortical ratio above 2. The corresponding step by step annotated codes will be available upon publication on our Reymann lab Github.

### Image analysis (extracts)

Quantification is performed using Fiji, Excel and RStudio. The last frame of the image sequence is used to define a region of interest manually, region in which the integrated raw intensity is followed over time. This ROI should include most of the actin filaments polymerised (Figure 5B, S5A-C) while excluding the cell/embryo ghost. The success of the cell lysis is assessed and experiments for which cells did not burst properly are excluded from the analysis (flow of material out the cell is visible in phase contrast after ablation). Measurement of the integrated fluorescence intensity is performed for each time frame and normalised using the first 10 frames, before averaging over the different embryos. The *C. elegans* embryonic cell volume and surface are taken from published publication (Thiels et al., 2021).

## Acknowledgements

The authors would like to warmly thank Stephan Grill and Sophie Quintin for critical comments on the manuscript. A special thanks for providing *C. elegans* strains to Karin Crell and Stephan Grill, notably for generating SWG300 strain as replacement of SWG004, as well as for SWG052; as well as Pierre Gönczy and Guangshuo Ou. Some strains were also provided by the CGC, which is funded by NIH Office of Research Infrastructure Programs (P40 OD010440). We warmly thank Christophe Guerin and Laurent Blanchoin for providing actin and actin binding proteins as well as for training us for microcavities fabrication. We would like also to particularly thank the Imaging Center of IGBMC (ici.igbmc.fr) for their assistance. We are also grateful to additional members of A-C.R. laboratory for discussion and technical assistance, as well as Christophe Reymann for insightful discussions on quantification methods.

---

**Figure S1:**
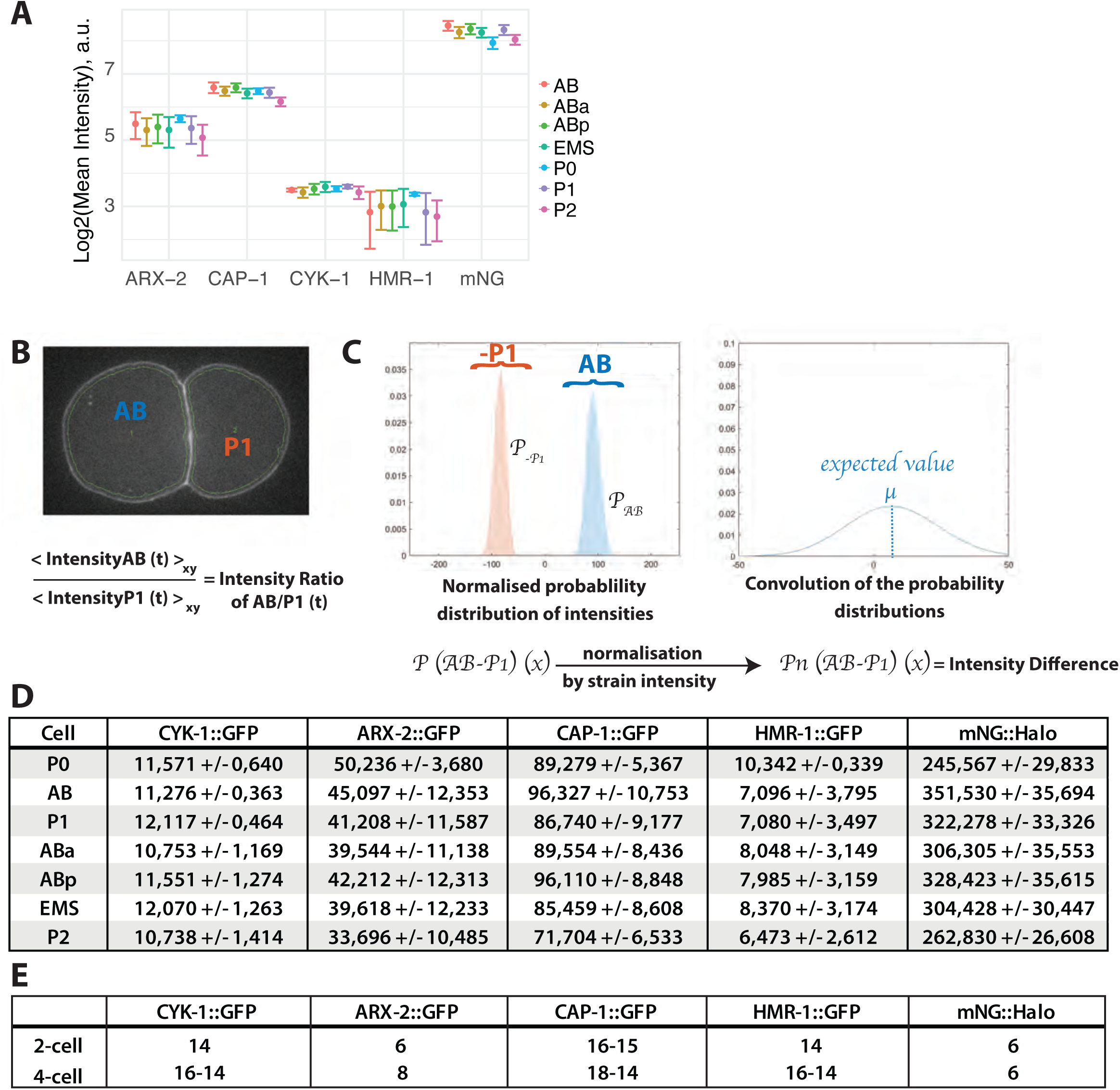
(A) Overall mean fluorescent intensity in each cell measured in the cytoplasm (mean ± standard deviation), GFP intensities for all actin related proteins acquired under the same imaging conditions. (B) Example of a cell segmentation output from Illastik. Cell ROIs are used to extract intensities such as the ratio between AB and P1 cell pair at each time point. (C) Example of plots of normalised probability distribution of the intensities of these two cells (AB and -P1 are shown) as well as the convolution of these probability distributions (AB-P1). The extracted expected value and standard deviation of these probability distributions are referred to as cell-cell intensity difference in the main text. (D) Table of average fluorescence intensities for each cell measured in the cytoplasm (mean ± standard deviation) and used to normalise the probability distribution. (E) Table of the number of cells counted for each category. Note that some cells are removed for the analysis such as if the acquired and segmented cortical surface is two small (cortex out of plane).

**Figure S2:**
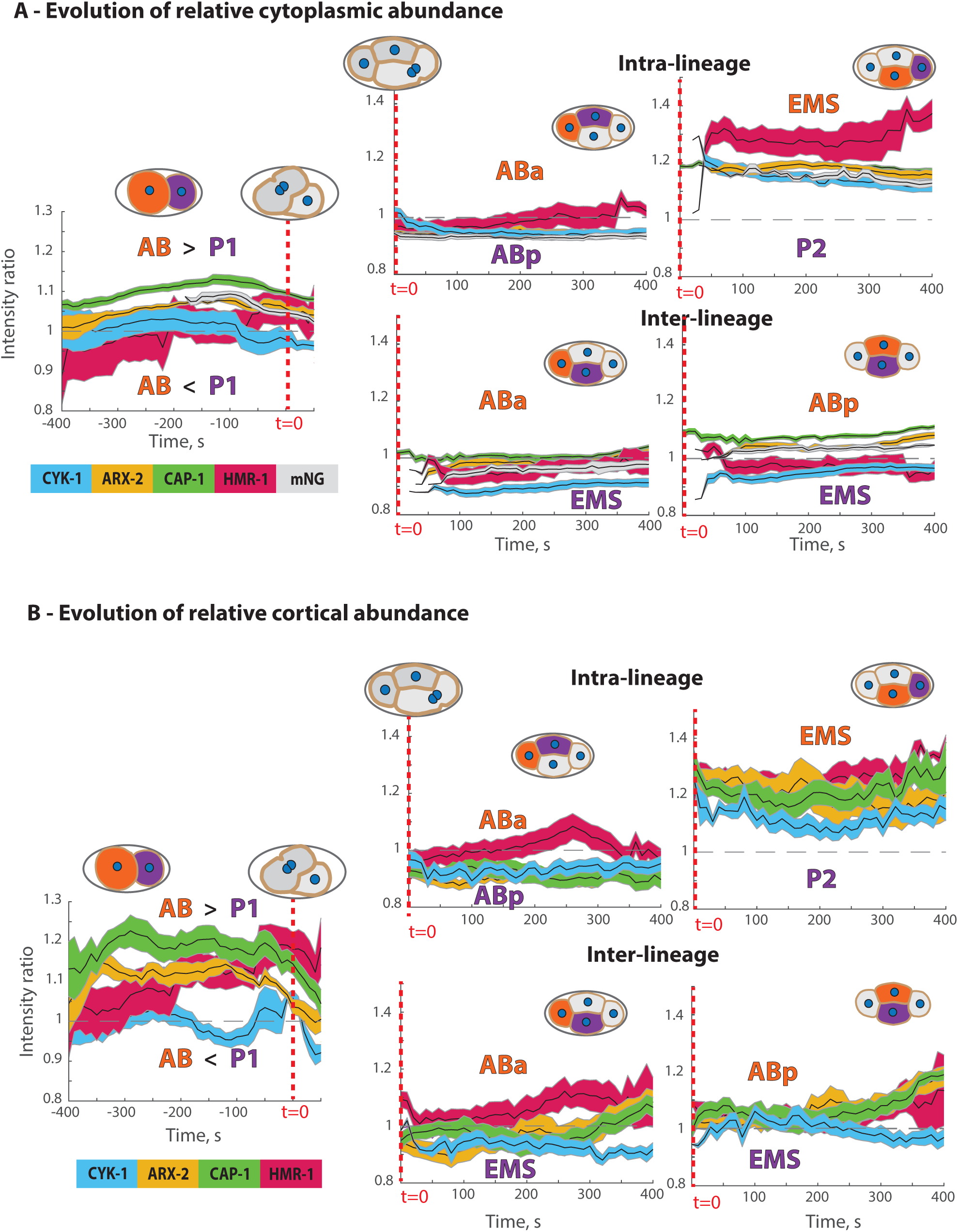
Time evolution of relative protein abundances. Time evolution of the intensity ratio for the different cell pairs in the cytoplasm (A) and in the cortical plane (B). Time is aligned using the reference timing illustrated in Figure 1A (red dashed lines).

**Figure S3:**
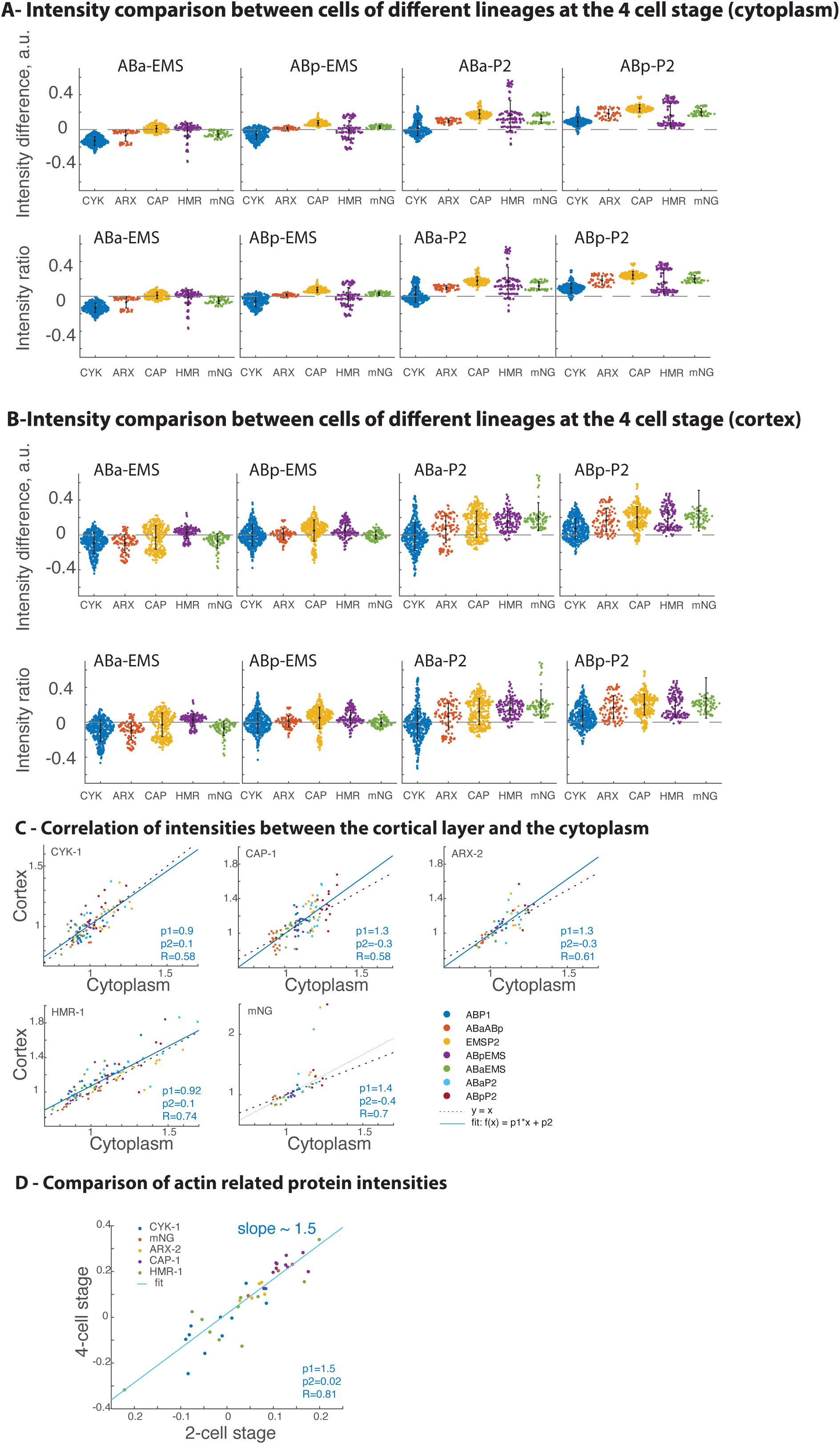
(A-B) Relative cortical abundance of additional cell pairs of the 4-cell stage: ratio of intensities and differences of intensities in the cytoplasm (A) and in the cortical plane (B). (C) Correlation plot of intensities in the cortex and cytoplasm (all cells and proteins shown separately). Linear model from the fit is shown as continuous line (f(x) = p1*x+p2) and f(x)=x is shown as dashed lines. (D) Comparison of cytoplasmic intensities between the 2-cell state and 4-cell state (calculated using intensity differences between all cell pairs).

**Figure S4:**
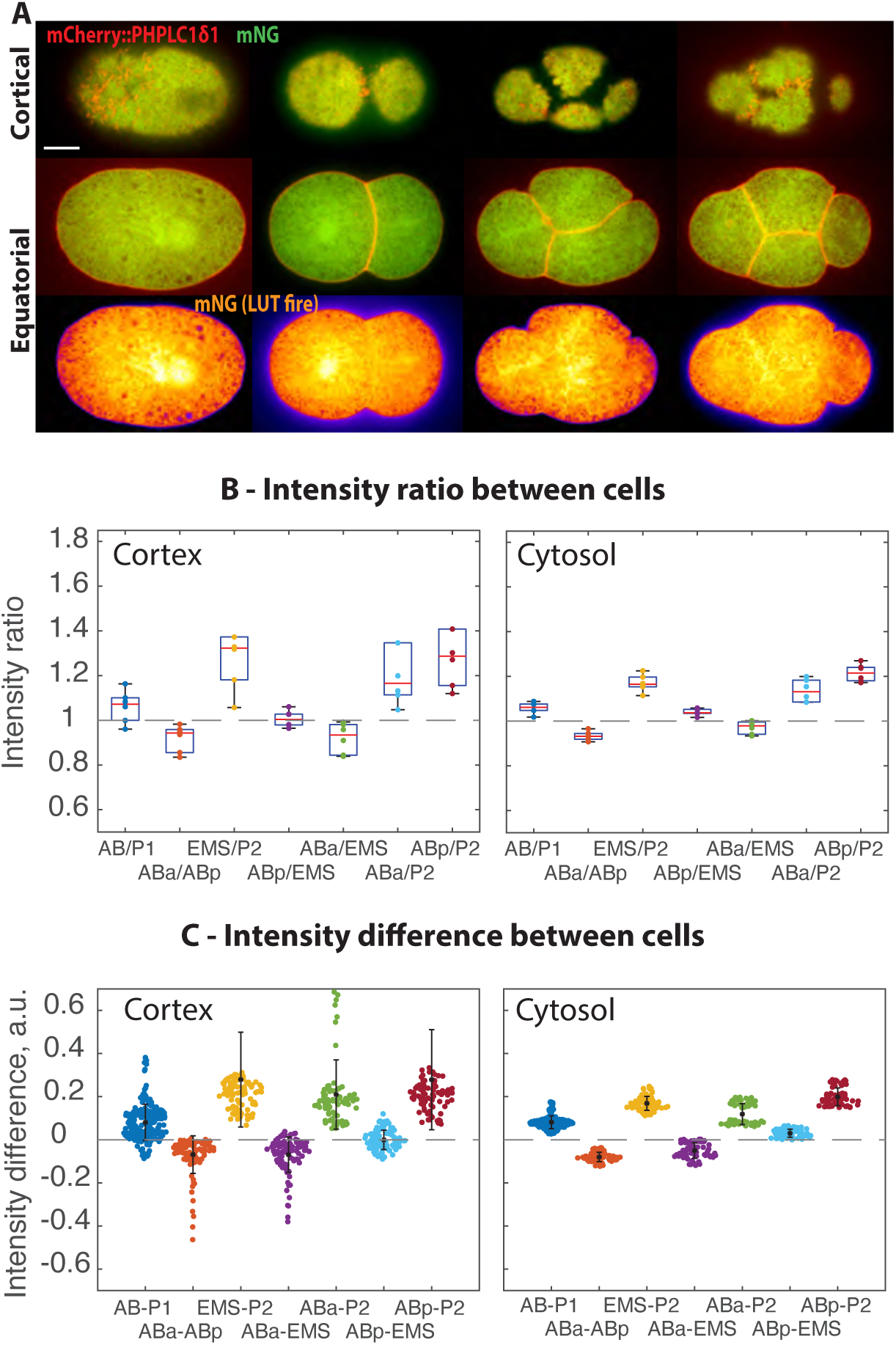
(A) Spatio-temporal distribution of freely diffusing mNG (green or fire) together with the membrane marker (PHPLC1δ1, red), at the cortex and equatorial plane of the embryo. Scale bar is 10 μm. (B) Ratio of intensities for each cell pair. (C) Differences of intensities for each cell pair, calculated via the convolution of normalised density probability distributions taken at each time point and normalised by the mean embryo intensity. Individual data points, mean values and standard deviations are shown for B and C.

**Figure S5:**
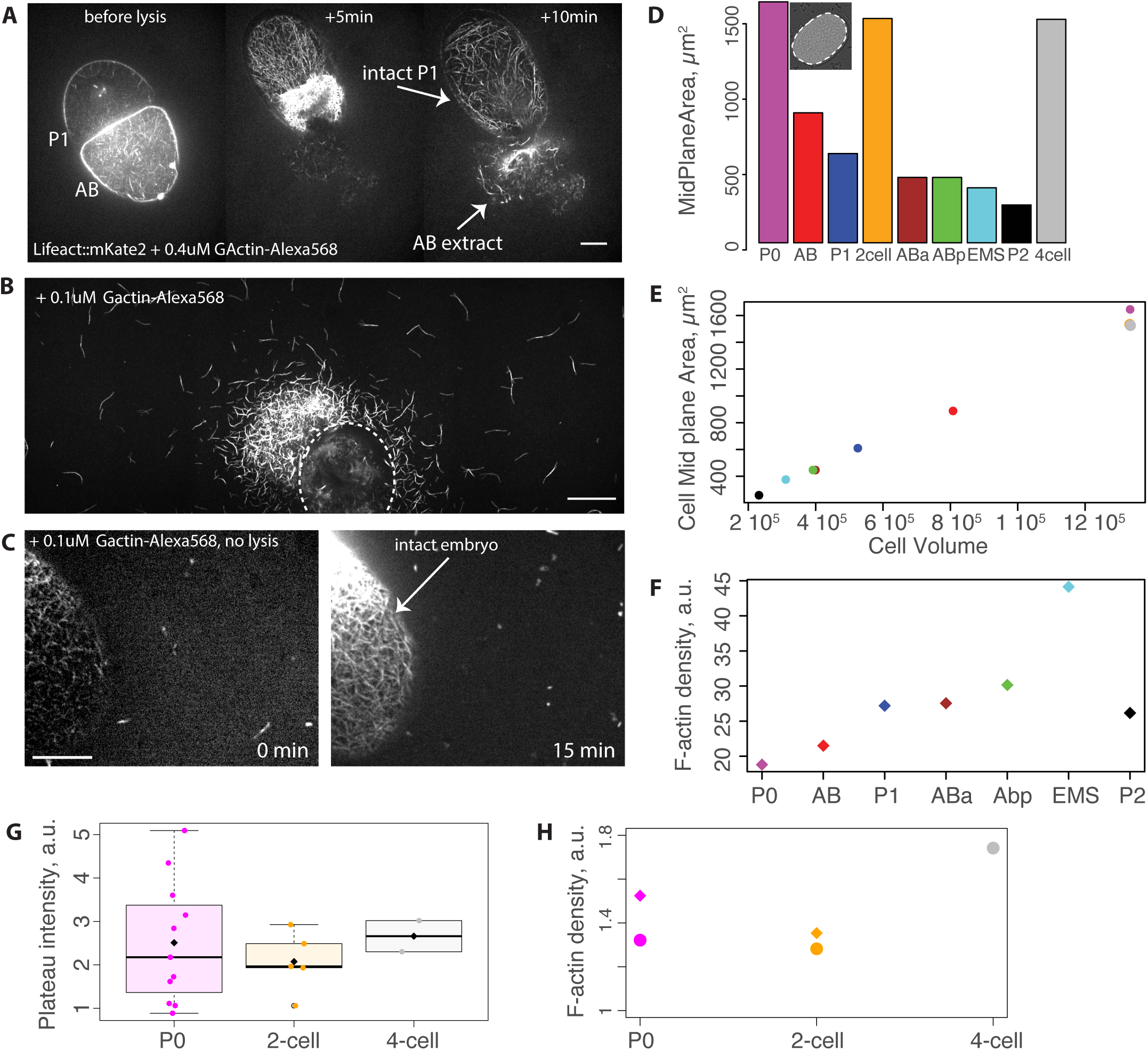
(A) Images of a 2-cell stage embryo before and after laser ablation, illustrating the ablation and extrusion of AB cell content. The intact P1 cell fills the entire eggshell space as AB extract spills out in the surrounding area. (B) Large field of view of a representative cell-extract polymerised in presence of 0.1μM G-actin-Alexa568 after UV laser ablation. The remaining embryo ghost (dashed circle) is out of plane- as acquisition is performed close to the coverslip in the plane where most of the filaments are polymerising due to the presence of methylcellulose. (C) Images of an embryo in absence of laser ablation in the same actin polymerisation media. Phalloidin penetrates the eggshell and membrane to label actin filaments within the cell cortex. (D) Quantification of midplane area from phase contrast images prior ablation. (E) Correlation between experimental mid plane cell area and published cell volumes. (F) F-actin density is calculated using the mean plateau intensity value normalised by published volumed of the corresponding cells, mean (diamond) and median (circle) are shown. (G) Plateau intensity values of P0 zygote as well as lysis at the 2-cell stage of AB+P1 cells and all 4 cells of the 4-cell stage. (H) F-actin density is calculated using the plateau intensity value normalised by the midplane area of the corresponding cells, mean (diamond) and median (circle) are shown. Scale bar: 20 μm for A and B, 10 μm for C.

## References

Ai, E., Skop, A.R., 2009. Endosomal recycling regulation during cytokinesis. Commun. Integr. Biol. 2, 444–447. 10.4161/cib.2.5.8931

Aroush, D.R.-B., Ofer, N., Abu Shah, E., Allard, J., Krichevsky, O., Mogilner, A., Keren, K., 2017. Actin Turnover in Lamellipodial Fragments. Curr. Biol. 27, 2963–2973.e14.

Audhya, A., Desai, A., Oegema, K., 2007. A role for Rab5 in structuring the endoplasmic reticulum. J. Cell Biol. 178, 43–56. 10.1083/jcb.200701139

Beatty, A., Morton, D.G., Kemphues, K., 2013. PAR-2, LGL-1 and the CDC-42 GAP CHIN-1 act in distinct pathways to maintain polarity in the C. elegans embryo. Development 140, 2005– 2014. 10.1242/dev.088310

Berg, S., Kutra, D., Kroeger, T., Straehle, C.N., Kausler, B.X., Haubold, C., Schiegg, M., Ales, J., Beier, T., Rudy, M., Eren, K., Cervantes, J.I., Xu, B., Beuttenmueller, F., Wolny, A., Zhang, C., Koethe, U., Hamprecht, F.A., Kreshuk, A., 2019. ilastik: interactive machine learning for (bio)image analysis. Nat. Methods 16, 1226–1232. 10.1038/s41592-019-0582-9

Cao, L.G., Wang, Y.L., 1990. Mechanism of the formation of contractile ring in dividing cultured animal cells. II. Cortical movement of microinjected actin filaments. J. Cell Biol. 111, 1905– 1911.

Carlier, M.F., 1998. Control of actin dynamics. Curr. Opin. Cell Biol. 10, 45–51. 10.1016/s0955-0674(98)80085-9

Caroti, F., Thiels, W., Vanslambrouck, M., Jelier, R., 2021. Wnt Signaling Induces Asymmetric Dynamics in the Actomyosin Cortex of the C. elegans Endomesodermal Precursor Cell. Front. Cell Dev. Biol. 9, 702741. 10.3389/fcell.2021.702741

Chan, F.-Y., Silva, A.M., Saramago, J., Pereira-Sousa, J., Brighton, H.E., Pereira, M., Oegema, K., Gassmann, R., Carvalho, A.X., 2018. The ARP2/3 complex prevents excessive formin activity during cytokinesis. Mol. Biol. Cell 30, 96–107. 10.1091/mbc.E18-07-0471

Costache, V., Prigent Garcia, S., Plancke, C.N., Li, J., Begnaud, S., Suman, S.K., Reymann, A.-C., Kim, T., Robin, F.B., 2022. Rapid assembly of a polar network architecture drives efficient actomyosin contractility. Cell Rep. 39, 110868. 10.1016/j.celrep.2022.110868

Courtemanche, N., Pollard, T.D., Chen, Q., 2016. Avoiding artefacts when counting polymerized actin in live cells with LifeAct fused to fluorescent proteins. Nat. Cell Biol. 18, 676–683.

Cramer, L.P., Briggs, L.J., Dawe, H.R., 2002. Use of fluorescently labelled deoxyribonuclease I to spatially measure G-actin levels in migrating and non-migrating cells. Cell Motil. Cytoskeleton 51, 27–38. 10.1002/cm.10013

Daniels, B.R., Dobrowsky, T.M., Perkins, E.M., Sun, S.X., Wirtz, D., 2010. MEX-5 enrichment in the C. elegans early embryo mediated by differential diffusion. Dev. Camb. Engl. 137, 2579–2585. 10.1242/dev.051326

Delattre, M., Goehring, N.W., 2021. Chapter Nine - The first steps in the life of a worm: Themes and variations in asymmetric division in C. elegans and other nematodes, in: Jarriault, S., Podbilewicz, B. (Eds.), Current Topics in Developmental Biology, Nematode Models of Development and Disease. Academic Press, pp. 269–308. 10.1016/bs.ctdb.2020.12.006

Dickinson, D.J., Schwager, F., Pintard, L., Gotta, M., Goldstein, B., 2017. A single-cell biochemistry approach reveals PAR complex dynamics during cell polarization. Dev. Cell 42, 416–434.e11. 10.1016/j.devcel.2017.07.024

Evans, T.C., Crittenden, S.L., Kodoyianni, V., Kimble, J., 1994. Translational control of maternal glp-1 mRNA establishes an asymmetry in the C. elegans embryo. Cell 77, 183–194. 10.1016/0092-8674(94)90311-5

Funk, J., Merino, F., Venkova, L., Heydenreich, L., Kierfeld, J., Vargas, P., Raunser, S., Piel, M., Bieling, P., 2019. Profilin and formin constitute a pacemaker system for robust actin filament growth. eLife 8, 1826.

Gan, W.J., Motegi, F., 2021. Mechanochemical Control of Symmetry Breaking in the Caenorhabditis elegans Zygote. Front. Cell Dev. Biol. 8.

Gilden, J., Krummel, M.F., 2010. Control of cortical rigidity by the cytoskeleton: Emerging roles for septins. Cytoskeleton 67, 477–486. 10.1002/cm.20461

Goehring, N.W., Grill, S.W., 2013. Cell polarity: mechanochemical patterning. Trends Cell Biol. 23, 72–80. 10.1016/j.tcb.2012.10.009

Gönczy, P., Rose, L.S., 2005. Asymmetric cell division and axis formation in the embryo. WormBook Online Rev. C Elegans Biol. 1–20. 10.1895/wormbook.1.30.1

Gonzalez Rodriguez, S., Wirshing, A.C.E., Goodman, A.L., Goode, B.L., 2023. Cytosolic concentrations of actin binding proteins and the implications for in vivo F-actin turnover. J. Cell Biol. 222, e202306036. 10.1083/jcb.202306036

Griffin, E.E., Odde, D.J., Seydoux, G., 2011. Regulation of the MEX-5 gradient by a spatially segregated kinase/phosphatase cycle. Cell 146, 955–968. 10.1016/j.cell.2011.08.012

Hirani, N., Illukkumbura, R., Bland, T., Mathonnet, G., Suhner, D., Reymann, A.-C., Goehring, N.W., 2019. Anterior-enriched filopodia create the appearance of asymmetric membrane microdomains in polarizing C. elegans zygotes. J. Cell Sci. 132. 10.1242/jcs.230714

Hoege, C., Hyman, A.A., 2013. Principles of PAR polarity in Caenorhabditis elegans embryos. Nat. Rev. Mol. Cell Biol. 14, 315–322.

Horvitz, H.R., Herskowitz, I., 1992. Mechanisms of asymmetric cell division: two Bs or not two Bs, that is the question. Cell 68, 237–255. 10.1016/0092-8674(92)90468-r

Jordan, S.N., Davies, T., Zhuravlev, Y., Dumont, J., Shirasu-Hiza, M., Canman, J.C., 2016. Cortical PAR polarity proteins promote robust cytokinesis during asymmetric cell division. J. Cell Biol. 212, 39–49. 10.1083/jcb.201510063

Kiuchi, T., Ohashi, K., Kurita, S., Mizuno, K., 2007. Cofilin promotes stimulus-induced lamellipodium formation by generating an abundant supply of actin monomers. J. Cell Biol. 177, 465–476. 10.1083/jcb.200610005

Knight, J.K., Wood, W.B., 1998. Gastrulation initiation in Caenorhabditis elegans requires the function of gad-1, which encodes a protein with WD repeats. Dev. Biol. 198, 253–265. 10.1016/S0012-1606(98)80003-1

Koestler, S.A., Rottner, K., Lai, F., Block, J., Vinzenz, M., Small, J.V., 2009. F- and G-actin concentrations in lamellipodia of moving cells. PloS One 4, e4810. 10.1371/journal.pone.0004810

Lee, C.W., Vitriol, E.A., Shim, S., Wise, A.L., Velayutham, R.P., Zheng, J.Q., 2013. Dynamic localization of G-actin during membrane protrusion in neuronal motility. Curr. Biol. CB 23, 1046–1056. 10.1016/j.cub.2013.04.057

Malla, M., Pollard, T.D., Chen, Q., 2022. Counting actin in contractile rings reveals novel contributions of cofilin and type II myosins to fission yeast cytokinesis. Mol. Biol. Cell 33, ar51. 10.1091/mbc.E21-08-0376

Motegi, F., Sugimoto, A., 2006. Sequential functioning of the ECT-2 RhoGEF, RHO-1 and CDC-42 establishes cell polarity in Caenorhabditis elegans embryos. Nat. Cell Biol. 8, 978–985.

Munro, E., Nance, J., Priess, J.R., 2004. Cortical Flows Powered by Asymmetrical Contraction Transport PAR Proteins to Establish and Maintain Anterior-Posterior Polarity in the Early <i> C. elegans< \ldots. Dev. Cell 7, 413–424.

Naganathan, S.R., Fürthauer, S., Rodriguez, J., Fievet, B.T., Jülicher, F., Ahringer, J., Cannistraci, C.V., Grill, S.W., 2018. Morphogenetic degeneracies in the actomyosin cortex. eLife 7, e37677. 10.7554/eLife.37677

Najafabadi, F.R., Leaver, M., Grill, S.W., 2022. Orchestrating nonmuscle myosin II filament assembly at the onset of cytokinesis. Mol. Biol. Cell 33, ar74. 10.1091/mbc.E21-12-0599

Nakayama, Y., Shivas, J.M., Poole, D.S., Squirrell, J.M., Kulkoski, J.M., Schleede, J.B., Skop, A.R., 2009. Dynamin Participates in the Maintenance of Anterior Polarity in the Caenorhabditis elegans Embryo. Dev. Cell 16, 889–900.

Nguyen, T.Q., Sawa, H., Okano, H., White, J.G., 2000. The C. elegans septin genes, unc-59 and unc-61, are required for normal postembryonic cytokineses and morphogenesis but have no essential function in embryogenesis. J. Cell Sci. 113, 3825–3837. 10.1242/jcs.113.21.3825

Pacquelet, A., 2017. Asymmetric Cell Division in the One-Cell C. elegans Embryo: Multiple Steps to Generate Cell Size Asymmetry. Results Probl. Cell Differ. 61, 115–140. 10.1007/978-3-319-53150-2_5

Padmanabhan, A., Ong, H.T., Zaidel-Bar, R., 2017. Non-junctional E-Cadherin Clusters Regulate the Actomyosin Cortex in the C. elegans Zygote. Curr. Biol. 27, 103–112. 10.1016/j.cub.2016.10.032

Pimpale, L.G., Middelkoop, T.C., Mietke, A., Grill, S.W., 2020. Cell lineage-dependent chiral actomyosin flows drive cellular rearrangements in early Caenorhabditis elegans development. eLife 9, e54930. 10.7554/eLife.54930

Pohl, C., Bao, Z., 2010. Chiral forces organize left-right patterning in C. elegans by uncoupling midline and anteroposterior axis. Dev. Cell 19, 402–412. 10.1016/j.devcel.2010.08.014

Priess, J.R., 2005. Notch signaling in the C. elegans embryo, in: WormBook: The Online Review of C. Elegans Biology [Internet]. WormBook.

Priess, J.R., Thomson, J.N., 1987. Cellular interactions in early C. elegans embryos. Cell 48, 241–250. 10.1016/0092-8674(87)90427-2

Raja Venkatesh, A., Le, K.H., Weld, D.M., Brandman, O., 2024. Diffusive lensing as a mechanism of intracellular transport and compartmentalization. eLife 12, RP89794. 10.7554/eLife.89794

Ray, S., Zaidel bar, 2021. Actin capping protein regulates actomyosin contractility to maintain germline architecture in C. elegans 19.

Reymann, A.-C., Staniscia, F., Erzberger, A., Salbreux, G., Grill, S.W., 2016. Cortical flow aligns actin filaments to form a furrow. eLife 5, e17807. 10.7554/eLife.17807

Robin, F.B., McFadden, W.M., Yao, B., Munro, E.M., 2014. Single-molecule analysis of cell surface dynamics in Caenorhabditis elegans embryos. Nat. Methods 11, 677–682.

Rocheleau, C.E., Downs, W.D., Lin, R., Wittmann, C., Bei, Y., Cha, Y.-H., Ali, M., Priess, J.R., Mello, C.C., 1997. Wnt Signaling and an APC-Related Gene Specify Endoderm in Early C. elegans Embryos. Cell 90, 707–716. 10.1016/S0092-8674(00)80531-0

Roh-Johnson, M., Goldstein, B., 2009. In vivo roles for Arp2/3 in cortical actin organization during C. elegans gastrulation. J. Cell Sci. 122, 3983–3993.

Roh-Johnson, M., Shemer, G., Higgins, C.D., McClellan, J.H., Werts, A.D., Tulu, U.S., Gao, L., Betzig, E., Kiehart, D.P., Goldstein, B., 2012. Triggering a Cell Shape Change by Exploiting Preexisting Actomyosin Contractions. Science 335, 1232–1235.

Rose, L., Gönczy, P., 2014. Polarity establishment, asymmetric division and segregation of fate determinants in early C. elegans embryos. WormBook 1–43.

Samandar Eweis, D., Plastino, J., 2020. Roles of Actin in the Morphogenesis of the Early Caenorhabditis elegans Embryo. Int. J. Mol. Sci. 21, 3652. 10.3390/ijms21103652

Scholze, M.J., Barbieux, K.S., De Simone, A., Boumasmoud, M., Süess, C.C.N., Wang, R., Gönczy, P., 2018. PI(4,5)P2 forms dynamic cortical structures and directs actin distribution as well as polarity in Caenorhabditis elegans embryos. Dev. Camb. Engl. 145, dev164988. 10.1242/dev.164988

Schonegg, S., Hyman, A.A., 2006. CDC-42 and RHO-1 coordinate acto-myosin contractility and PAR protein localization during polarity establishment in C. elegans embryos. Dev. Camb. Engl. 133, 3507–3516. 10.1242/dev.02527

Severson, A.F., Baillie, D.L., Bowerman, B., 2002. A Formin Homology Protein and a Profilin Are Required for Cytokinesis and Arp2/3-Independent Assembly of Cortical Microfilaments in C. elegans. Curr. Biol. 12, 2066–2075. 10.1016/S0960-9822(02)01355-6

Shivas, J.M., Skop, A.R., 2012. Arp2/3 mediates early endosome dynamics necessary for the maintenance of PAR asymmetry in Caenorhabditis elegans. Mol. Biol. Cell 23, 1917–1927. 10.1091/mbc.E12-01-0006

Shukla, Y., Ghatpande, V., Hu, C.F., Dickinson, D.J., Cenik, C., 2025. Landscape and regulation of mRNA translation in the early C. elegans embryo. Cell Rep. 44, 115778. 10.1016/j.celrep.2025.115778

Skruber, K., Read, T.-A., Vitriol, E.A., 2018. Reconsidering an active role for G-actin in cytoskeletal regulation. J. Cell Sci. 131, jcs203760-11.

Suarez, C., Kovar, D.R., 2016. Internetwork competition for monomers governs actin cytoskeleton organization. Nat. Rev. Mol. Cell Biol.

Sulston, J.E., Schierenberg, E., White, J.G., Thomson, J.N., 1983. The embryonic cell lineage of the nematode Caenorhabditis elegans. Dev. Biol. 100, 64–119. 10.1016/0012-1606(83)90201-4

Swan, K.A., Severson, A.F., Carter, J.C., Martin, P.R., Schnabel, H., Schnabel, R., Bowerman, B., 1998. cyk-1: a C. elegans FH gene required for a late step in embryonic cytokinesis. J. Cell Sci. 111, 2017–2027. 10.1242/jcs.111.14.2017

Tax, F.E., Thomas, J.H., 1994. Cell-Cell Interactions: Receiving signals in the nematode embryo. Curr. Biol. 4, 914–916. 10.1016/S0960-9822(00)00203-7

Tenlen, J.R., Molk, J.N., London, N., Page, B.D., Priess, J.R., 2008. MEX-5 asymmetry in one-cell C. elegans embryos requires PAR-4- and PAR-1-dependent phosphorylation. Dev. Camb. Engl. 135, 3665–3675. 10.1242/dev.027060

Thiels, W., Smeets, B., Cuvelier, M., Caroti, F., Jelier, R., 2021. spheresDT/Mpacts-PiCS: cell tracking and shape retrieval in membrane-labeled embryos. Bioinformatics 37, 4851–4856. 10.1093/bioinformatics/btab557

Velarde, N., Gunsalus, K.C., Piano, F., 2007. Diverse roles of actin in C. elegans early embryogenesis. BMC Dev. Biol. 7, 142. 10.1186/1471-213X-7-142

Vitriol, E.A., McMillen, L.M., Kapustina, M., Gomez, S.M., Vavylonis, D., Zheng, J.Q., 2015. Two functionally distinct sources of actin monomers supply the leading edge of lamellipodia. Cell Rep. 11, 433–445. 10.1016/j.celrep.2015.03.033

Wu, Y., Zhang, H., Griffin, E.E., 2015. Coupling between cytoplasmic concentration gradients through local control of protein mobility in the Caenorhabditis elegans zygote. Mol. Biol. Cell 26, 2963–2970. 10.1091/mbc.E15-05-0302

Yamamoto, K., Ichbiah, S., Perez, M., Borrego-Pinto, J., Delbary, F., Goehring, N.W., Northrop, P., Turlier, H., Charras, G., 2026. Spatiotemporal mapping of the contractile and adhesive forces sculpting early *C. elegans* embryos. Dev. Cell 61, 1273–1288.e4. 10.1016/j.devcel.2026.04.013

Yan, V.T., Narayanan, A., Wiegand, T., Jülicher, F., Grill, S.W., 2022. A condensate dynamic instability orchestrates actomyosin cortex activation. Nature 609, 597–604. 10.1038/s41586-022-05084-3

Zaatri, A., Perry, J.A., Maddox, A.S., 2021. Septins and a formin have distinct functions in anaphase chiral cortical rotation in the Caenorhabditis elegans zygote. Mol. Biol. Cell 32, 1283–1292. 10.1091/mbc.E20-09-0576

Zacharias, A.L., Murray, J.I., 2016. Combinatorial decoding of the invariant C. elegansembryonic lineage in space and time. genesis 54, 182–197.

Zhu, Z., Chai, Y., Jiang, Y., Li, Wenjing, Hu, H., Li, Wei, Wu, J.-W., Wang, Z.-X., Huang, S., Ou, G., 2016. Functional Coordination of WAVE and WASP in C. elegans Neuroblast Migration. Dev. Cell 39, 224–238. 10.1016/j.devcel.2016.09.029

